# Neuron-astrocyte metabolic coupling facilitates spinal plasticity and maintenance of persistent pain

**DOI:** 10.1101/2022.12.03.518519

**Authors:** Sebastián Marty-Lombardi, Shiying Lu, Wojciech Ambroziak, Hagen Wende, Katrin Schrenk-Siemens, Anna A. DePaoli-Roach, Anna M. Hagenston, Anke Tappe-Theodor, Manuela Simonetti, Rohini Kuner, Thomas Fleming, Jan Siemens

## Abstract

Long-lasting pain stimuli can trigger maladaptive changes in the spinal cord, reminiscent of plasticity associated with memory formation. Metabolic coupling between astrocytes and neurons has been implicated in neuronal plasticity and memory formation in the CNS, but neither its involvement in pathological pain nor in spinal plasticity has been tested. Here, we report a novel form of neuroglia signaling involving spinal astrocytic glycogen dynamics triggered by persistent noxious stimulation via upregulation of the metabolic signaling molecule PTG exclusively in spinal astrocytes. PTG drove glycogen build-up in astrocytes, and blunting glycogen accumulation and turnover by *Ptg* gene deletion reduced pain-related behaviors and promoted faster recovery by shortening pain maintenance. Furthermore, mechanistic analyses revealed that glycogen dynamics is a critically required process for maintenance of pain by facilitating neuronal plasticity in spinal lamina 1 neurons. Finally, metabolic analysis indicated that glycolysis and lactate transfer between astrocytes and neurons fuels spinal neuron hyperexcitability.

Spinal glycogen-metabolic cascades therefore hold therapeutic potential to alleviate pathological pain.

## Introduction

Sensory stimuli, in particular those that are strong, long-lasting or repetitive, can trigger neuronal plasticity. These activity-dependent processes require metabolic energy to fuel ensuing structural and functional changes^1^. The resulting neurophysiological changes, which often involve alterations of synaptic properties and changing of synaptic strength, can be beneficial and allow, for example, adaptation to a changing environment. But persistent stimulation in a pathological context can also give rise to chronic pain, a maladaptive form of neuronal plasticity detrimental to health^2^.

The spinal cord is the first relay station of noxious signals, where primary afferent sensory neurons connect to projection neurons residing in the dorsal spinal cord that carry signals of potentially harmful and damaging (thus painful) stimuli to higher brain centers. A multitude of additional spinal excitatory and inhibitory neurons modulate and “gate” painful signals^3^. Plastic changes within this extensive neuronal network have long been associated with pathological forms of pain^4^. In parallel, many studies have emphasized the role of non-neuronal cells, such as astrocytes and microglia, in spinal nociceptive signal processing and pain chronification. Astrocytes provide metabolic support to neurons, regulate extracellular ion composition and modulate synaptic transmission, and are thus important for physiological (homeostatic) neuronal processing in the CNS^5^. Astrocytes have also been implicated in modulating nociceptive signaling along the spinothalamic axis and, in this context, they play a preeminent role in the dorsal spinal cord. In pathological contexts, spinal astrocytes have been found to not only modulate the induction of long-lasting pain but appear to be particularly relevant for the maintenance of chronic pain states^6–9^. How spinal astrocytes and neurons interact to drive pathological pain states and whether this interaction can be therapeutically harnessed for pain therapy are important medical questions.

Energy-carrying metabolites such as lactate can modulate nociceptive signal processing in spinal circuits^10^. However, the relevance and potential source of such metabolites in pathological nociceptive processing have remained largely unknown. Moreover, genetic evidence has been missing that directly links the regulation of energy metabolism and spinal nociceptive signal processing *in vivo*.

Here we report a novel metabolic mechanism at the interface between the peripheral and central nervous system that regulates excitability of spinal neurons processing and propagating noxious information. We find that spinal astrocytes and neurons are metabolically coupled and that strong, long-lasting pain stimuli have pronounced effects on dynamic glycogen metabolism in astrocytes. Robust and protracted glycogen build-up in spinal astrocytes is mediated by noxious stimulation-induced transcriptional activation of PTG (<u>P</u>rotein <u>T</u>argeting to <u>G</u>lycogen). Genetically perturbing glycogen accumulation and dynamics by deleting the *Ptg* gene from astrocytes does not affect short-term/acute detection of noxious, pain-inducing signals in mice but allows animals to recover faster from long-lasting inflammatory pain. Mechanistically, we find that spinal neuronal plasticity, induced by long-lasting inflammatory pain stimulation, requires astrocytic glycogen and energy metabolism. Our study demonstrates that astrocytic energy fuels the maintenance of long-lasting inflammatory pain states and that pharmacologic interference with spinal astrocyte-neuron energy coupling may constitute a therapeutic avenue to accelerate recovery from pathological forms of pain by inhibiting maladaptive neuronal plasticity.

## Results

### Noxious, pain-inducing stimulation triggers robust PTG expression in spinal astrocytes

We utilized a ribosomal profiling screen to identify mRNA transcripts actively translated in the spinal cord upon noxious stimulation that lead to pain-related behaviors in mice. This approach capitalizes on the finding that a structural component of the ribosome, the S6 protein, becomes phosphorylated––both in neuronal and non-neuronal cells––upon a wide variety of stimuli^11^. Using antibodies against the phospho-moiety of S6 (pS6), it is possible to biochemically isolate activated polysomes from tissue of stimulated animals and identify actively translated mRNAs by RNAseq^11–13^. Using anti-pS6 antibodies, we found that pain stimuli effectively induced S6 protein phosphorylation in the spinal cord of mice (**Extended Data Fig. 1a-d**). We tested several anti-pS6 antibodies and identified clone 5364 as the most effective antibody for biochemical isolation and enrichment of pain stimulation-activated polysomes from the mouse dorsal spinal cord (**Extended Data Fig. 1e,f**).

**Fig. 1:**
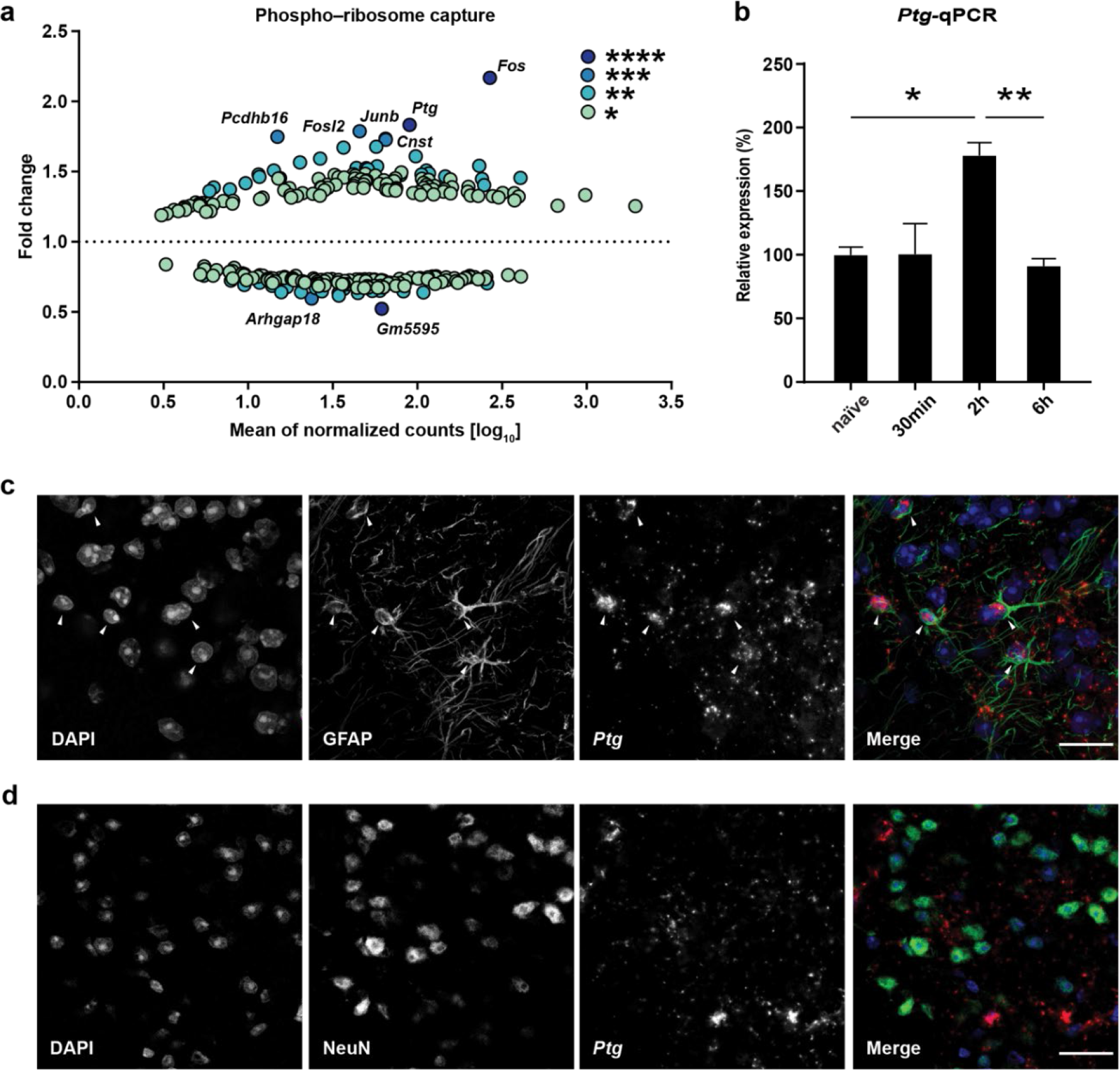
Noxious, pain-inducing stimulation triggers *Ptg* expression in astrocytes of the dorsal spinal cord. **a**, RNAseq of pS6 captured ribosomes isolated from ipsilateral and contralateral spinal dorsal horn 2 hours after mice had been subjected to formalin stimulation. Y-axis indicates relative enrichment or de-enrichment (ratio) of transcripts detected in ipsilateral (affected) dorsal horn tissue vs. contralateral tissue. X-axis indicates relative mRNA abundance. Only significantly changed transcripts are shown (color code indicates p-value, Deseq2, N=3). **b**, qPCR-determined relative expression level of the *Ptg* gene at different time points after formalin stimulation (N=3 mice). One-way ANOVA with Tukey’s post hoc test. **c**,**d**, Representative images of combined immunohistochemistry and fluorescent *in situ* hybridizations with antibodies directed against GFAP (**c**, green) and NeuN (**d**, green) and an *in situ* probe directed against *Ptg* (red), DAPI (blue); scale bar: 20µm; data represent mean ± s.e.m. *p < 0.05, **p < 0.01, ***p < 0.005, ****p < 0.001. See also Extended Data Fig. 1.

We expected that transcripts of the immediate early genes *cFos* and *FosB*, which are induced in the spinal cord upon pain stimulation^14^ (**Extended Data Fig. 1b**), would be enriched in biochemically isolated pS6 ribosome pull-downs, which was indeed the case (**Extended Data Fig. 1g**). Also, pS6 ribosomal profiling slightly but significantly enriched cFos transcripts compared to total lysates obtained from the spinal cord of pain-stimulated mice (**Extended Data Fig. 1h**), suggesting that pS6 ribosomal profiling is a slightly more sensitive method to detect actively transcribed-translated genes compared to simply using spinal cord tissue for direct transcriptional profiling of spinal cord lysates.

Injecting Formalin (FA) into the intraplantar surface of the mouse paw is a widely used pain model^15^. When ipsilateral (the FA-stimulated side of the spinal cord) and contralateral (the non-FA injected control side) spinal cord sections, corresponding to the relevant (hindpaw) dermatome levels L3-L6 (Extended Data Fig. 1a,c**,d**), were compared by pS6 ribosomal profiling, we identified *Ptg* (<u>P</u>rotein <u>T</u>argeting to <u>G</u>lycogen; also referred to as *Ppp1r3c*) as the most robustly induced gene, next to *cFos* (**Fig. 1a**). Pain-induced *Ptg* expression was verified by qPCR (**Fig. 1b**) and multi-color *in situ* hybridization (**Extended Data Fig. 1d**), confirming that *Ptg* induction is confined to the pain stimulation-affected (ipsilateral) side of the dorsal spinal cord.

On the cellular level, we found astrocytes—rather than neurons—express *Ptg* upon pain stimulation (**Fig. 1c,d**). Next to spinal astrocytes and neurons, also spinal microglia have been implicated in pathological forms of pain^16–18^. However, we did not find any *Ptg* expression in microglia in the context of noxious FA stimulation (**Extended Data Fig. 1i**).

### Spinal glycogen levels dynamically change subsequent to noxious stimulation

PTG is a regulator of glucose- and glycogen metabolism in that the molecule directs the protein phosphatase 1 (PP1) to glycogen synthase and glycogen phosphorylase to promote dephosphorylation of these two key anabolic/catabolic glycogen enzymes thereby inducing glycogen formation and inhibiting glycogen break-down, respectively^19^.

Of note, in the brain glycogen is primarily stored in astrocytes^20–22^ and *in vitro* studies of brain-derived neuron-astrocyte co-cultures show that forced expression of PTG in cultured astrocytes is sufficient to drive glycogen build-up^23^.

We therefore wondered whether a noxious stimulus that induces pain would result in increased spinal glycogen levels. Indeed, glycogen levels increased on the ipsilateral- but not on the contralateral side of the spinal cord subsequent to FA-induced pain (**Fig. 2a**). Interestingly, glycogen level increases were detectable with a delay of 6 hours after the stimulus (4 hours after *Ptg* mRNA peaked) and dropped to baseline levels within 1 to 3 days.

**Fig. 2:**
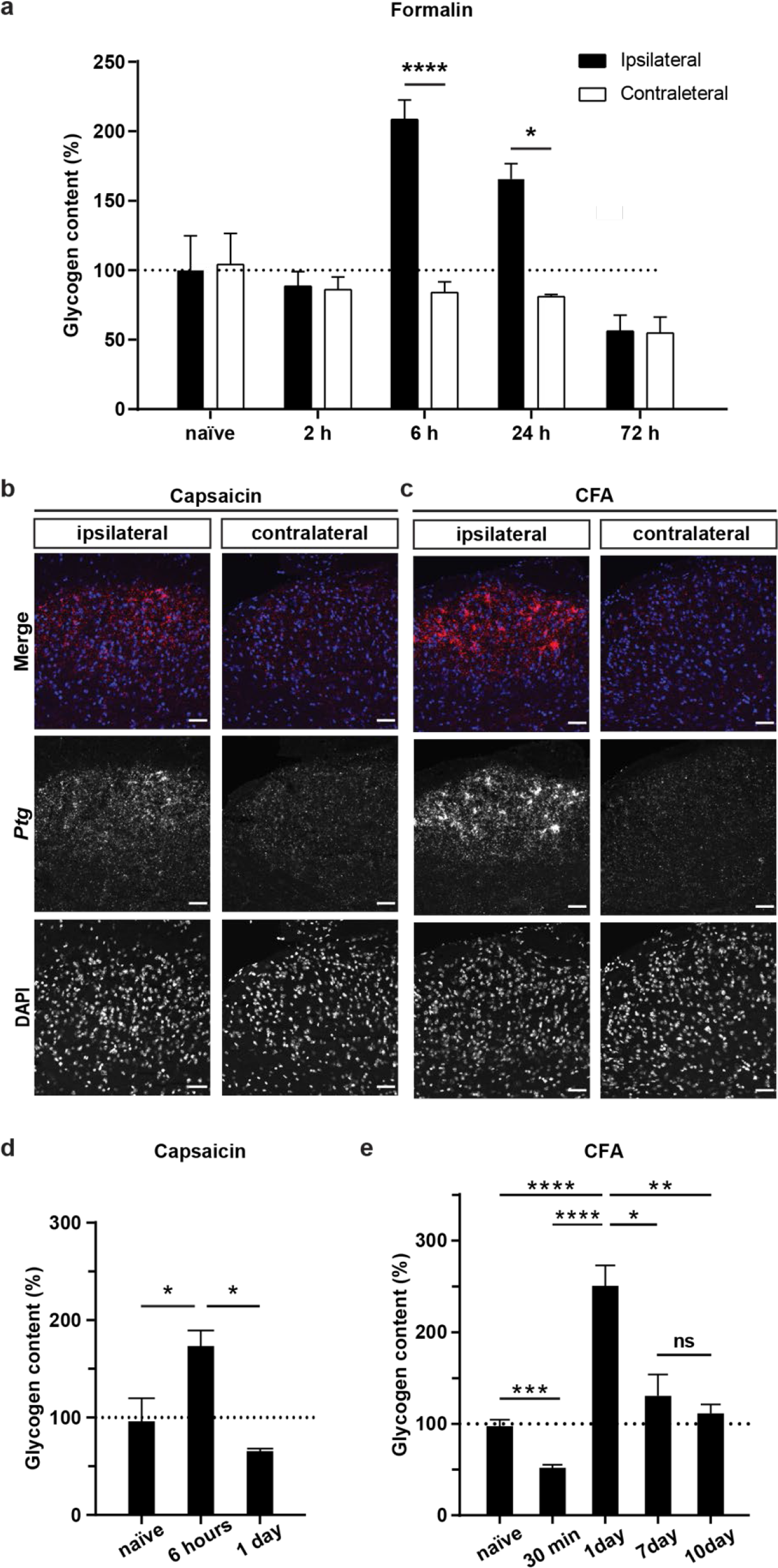
Inflammatory pain induces long-lasting glycogen accumulation in the dorsal spinal cord. **a**, Glycogen content of ipsilateral and contralateral dorsal spinal cord tissue isolated at different time points after formalin stimulation and expressed as percent of glycogen of ipsilateral tissue isolated from naïve mice (N=5); Two-way ANOVA with Bonferroni post hoc test. **b**,**c**, Representative images of *in situ* hybridizations (RNAscope) with a probe for *Ptg* (red) of spinal cord tissue 2 hours after capsaicin stimulation (**b**) or subjecting mice to the CFA inflammatory pain model (**c**); DAPI (blue), scale bar: 50µm. **d**,**e** Glycogen content (percent of naïve) of dorsal spinal cord tissue at different time points after capsaicin- (**d**) or CFA stimulation (**e**). (N=3); One-way ANOVA with Tukey’s post hoc test. Data represent mean ± s.e.m. *p < 0.05, **p < 0.01, ***p < 0.005, ****p < 0.001. See also Extended Data Fig. 2.

Next, we asked whether other types of painful stimuli would also trigger *Ptg* induction and protracted glycogen increases in the spinal cord. Similar to FA, also stimulation using capsaicin and Complete Freud’s Adjuvant (CFA) induced *Ptg* mRNA expression on the ipsilateral side of the dorsal spinal cord (**Fig. 2b,c**). When assessing glycogen levels, we found that the shorter-acting capsaicin also induced glycogen elevation, albeit less pronounced compared to that of FA (**Fig. 2d**). On the other hand, CFA, a potent inducer of inflammatory pain that persists longer than FA-induced pain^15, 24, 25^, triggered glycogen increases that outlasted those observed for the FA pain model (**Fig. 2e**). Additionally, when assessing glycogen levels shortly after a noxious, pain-inducing stimulus, we found that glycogen levels initially––and very transiently––decreased (**Fig. 2e**).

These results suggest that a stronger and longer-lasting noxious stimulus resulted in stronger and longer-lasting glycogen build-up subsequent to the acute phase of the pain-inducing stimulus. We therefore also tested whether the combination of two disparate painful stimuli would result in additive glycogen accumulation. Spared-nerve injury (SNI), a model of neuropathic pain^26^, by itself only slightly increased glycogen levels, similar to a short-term capsaicin stimulation (**Extended Data Fig. 2a**). However, combing the two pain models resulted in synergistically increased spinal glycogen accumulation compared to mice subjected to either SNL- or capsaicin treatment alone (**Extended Data Fig. 2a**).

Collectively, these results demonstrate that pronounced glycogen dynamics accompany diverse inflammatory pain stimuli in the spinal cord and appear to be graded, at least to some extent, by the intensity and duration of noxious pain-inducing stimulus.

From the dorsal spinal cord, pain signals are relayed for further processing to higher order brain centers. We wondered whether a pain-stimulated glycogen increase is specific to the spinal cord or whether other brain areas also show increased glycogen levels. CFA-induced pain did not alter glycogen levels in the somatosensory-, insula- or prefrontal cortices or in the amygdala (**Extended Data Fig. 2b-e**), brain regions all of which are implicated in pain signal processing. While we cannot exclude that small, cell-type selective glycogen changes also occur in other brain areas involved in pain signal processing, strong and long-lasting painful insults appear to primarily modulate glycogen dynamics in the spinal cord.

### Noxious stimuli-induced glycogen build-up is blunted in astrocyte-specific PTG knock-out mice

Because pain-induced glycogen increases followed Ptg mRNA expression with a delay of a few hours—allowing for the translation of PTG protein in due course—we reasoned that spinal glycogen build-up is dependent on PTG protein expression. We therefore generated conditional (floxed) PTG knock-out animals (**Extended Data Fig. 3a**). We first crossed these conditional PTG knock-out mice with a Cre deleter mouse strain^27^ to delete functional PTG from the mouse germline and all tissues to obtain complete (global) PTG knock-out animals (gPTG^(-/-)^). We verified that pain-stimulated *Ptg* induction in spinal astrocytes was abolished in gPTG^(-/-)^ mice using multi-color *in situ* hybridization (**Fig. 3a and Extended Data Fig. 3b**). Moreover, when subjecting gPTG^(^**^-/-)^** mice to CFA-induced pain, we found that the spinal glycogen increase was indeed completely blunted compared to wildtype control mice (**Fig. 3b**). Even baseline glycogen appeared to be reduced in gPTG^(-/-)^ mice to levels such as were observed shortly after pain stimulation in wildtype mice (**Fig. 3b and Extended Data Fig. 3c**). To assess whether these effects are mediated by astrocytic PTG, we next crossed floxed-PTG mice to Aldh1L1-Cre^ERT2^ mice^28^ to ablate functional PTG from astrocytes by tamoxifen injections (we refer to these astrocytically *Ptg* deleted mice from here on as cPTG^(-/-)^ mice). Again, we confirmed absence of *Ptg* transcripts from spinal astrocytes of conditionally mutant animals (**Fig. 3a and Extended Data Fig. 3b**). Similar to gPTG^(-/-)^ mice, in these astrocytically deleted cPTG^(-/-)^ mice, CFA pain-induced glycogen build- up was also completely blunted (**Fig. 3c**). Different from gPTG^(-/-)^ mice, however, baseline glycogen levels appeared normal in cPTG^(-/-)^ animals (**Fig. 3c and Extended Data Fig. 3c**), suggesting that non-astrocytic (yet PTG-dependent) glycogen stores may contribute to baseline spinal glycogen levels. Alternatively, it is also possible that the conditional knock-out approach did not affect baseline astrocyte glycogen levels within the timeframe of the tamoxifen-induced PTG deletion (starting 3 weeks prior to the experiment), because glycogen levels might be fairly stable in the absence of any painful stimulus. Regardless of the origin of the low baseline glycogen levels, very clearly astrocytic PTG is required for nociceptive glycogen dynamics.

**Fig. 3:**
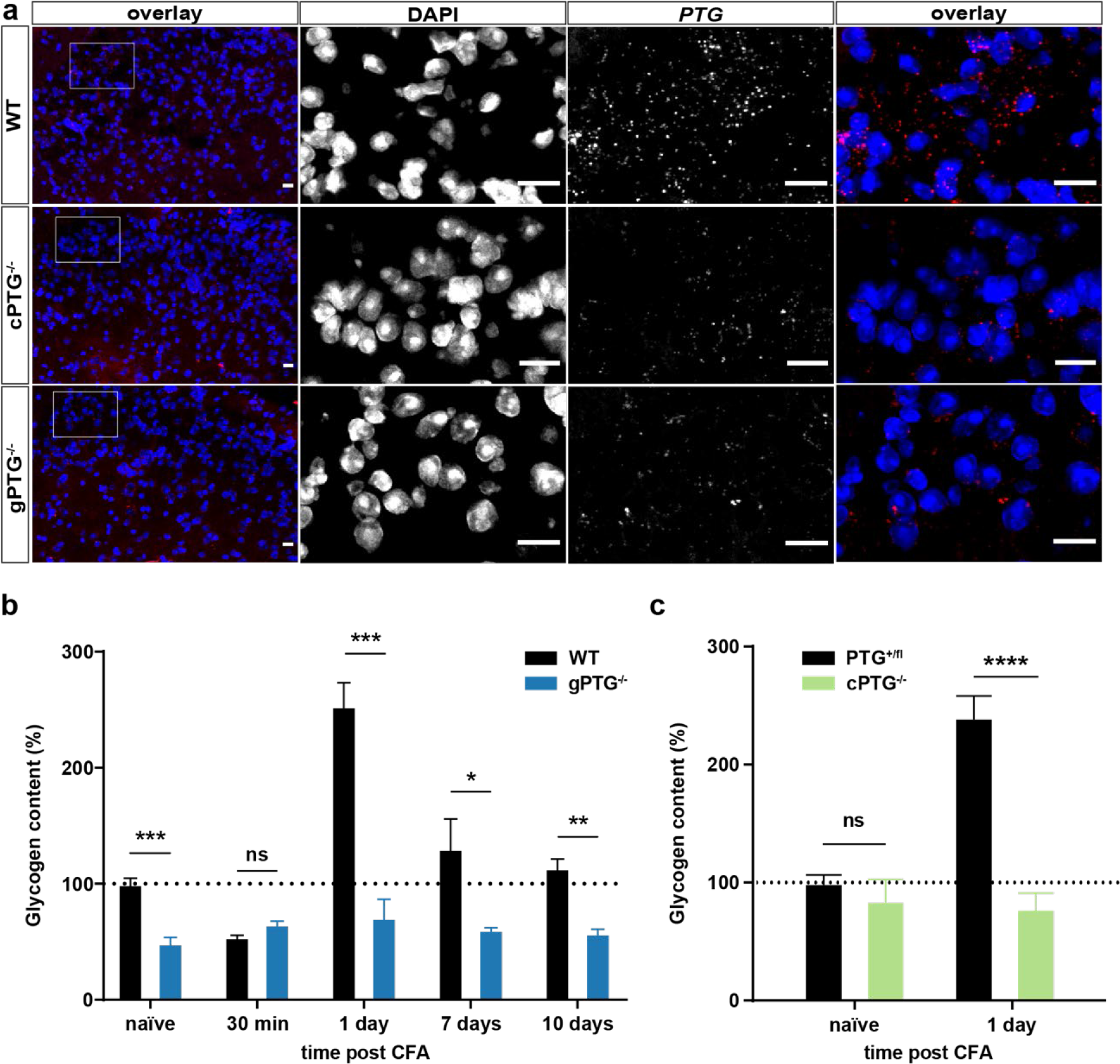
Noxious stimulation-induced spinal glycogen dynamics is blunted in PTG^(-/-)^ mice. **a**, Representative images of *in situ* hybridizations (RNAscope) with a probe for *Ptg* (red) of ipsilateral dorsal spinal cord tissue from wildtype, cPTG^(-/-)^ and gPTG^(-/-)^ mice 2 hours after formalin-induced pain. DAPI (blue); scale bar: 20 µm. **b**,**c**, Glycogen content (percent of naïve) of dorsal spinal cord tissue at different time points after CFA stimulation for gPTG^(-/-)^ (**b**, N=5) or cPTG^(-/-)^ (**c**, N=5) mice compared to their respective control mice. Two-way ANOVA with Bonferroni post hoc test. Data represent mean ± s.e.m. *p < 0.05, **p < 0.01, ***p < 0.005, ****p < 0.001. See also Extended Data Fig. 3.

### PTG deletion promotes faster recovery from long-lasting pain-related behavior

Given the dependence of noxious stimuli-induced spinal glycogen metabolism on PTG, we wondered if perturbation of astrocytic glycogen dynamics by deleting PTG would have any effect on behavioral pain-related responses in mice. In assessing pain-related behaviors, the measurement of the threshold and latency to (painful) mechanical and thermal stimuli applied to the animal’s hindpaw are standard pain testing paradigms^29^. Accordingly, we found that, under basal conditions, mechanical stimuli generated by poking the plantar hindpaw surface using von Frey filaments of different mechanical strength evoked normal responses in gPTG^(-/-)^ mice compared to control animals (**Fig. 4a,b**). Similarly, PTG deletion did not affect the latency of heat-pain triggered paw withdrawal when a light beam was focused on the plantar surface of the hindpaw (**Extended Data Fig. 4a**). We next evaluated whether PTG deletion would influence short-term pain sensitization such as that triggered by the intraplantar injection of capsaicin, serotonin, prostaglandin E2 (PGE2), or FA. We found that capsaicin provoked similar mechanical hypersensitivity in wildtype and cPTG^(-/-)^ mice (**Extended Data Fig. 4b**). Only inflammatory stimuli mediating longer-lasting pain sensitization, such as serotonin or prostaglandin E2 (PGE2)^30^, were associated with reduced mechanical hypersensitivity in gPTG^(-/-)^ mice (**Fig. 4c,d**). FA injection, which evokes spontaneous nocifensive responses such as licking, flinching and guarding of the affected paw, produced a slightly smaller response in the gPTG^(-/-)^ mice in the initial phase of the biphasic pain response. This difference appeared to be absent from cPTG^(-/-)^ mice, suggesting that intact baseline glycogen levels may be relevant for the early acute phase of FA-evoked nocifensive pain behavior (**Fig. 4e,f and Extended Data Fig. 4c,d**). Importantly, the more pronounced second phase of the FA-triggered response was indistinguishable between wildtype and both KO models (**Fig. 4e,f and Extended Data Fig. 4c,d**).

**Fig. 4:**
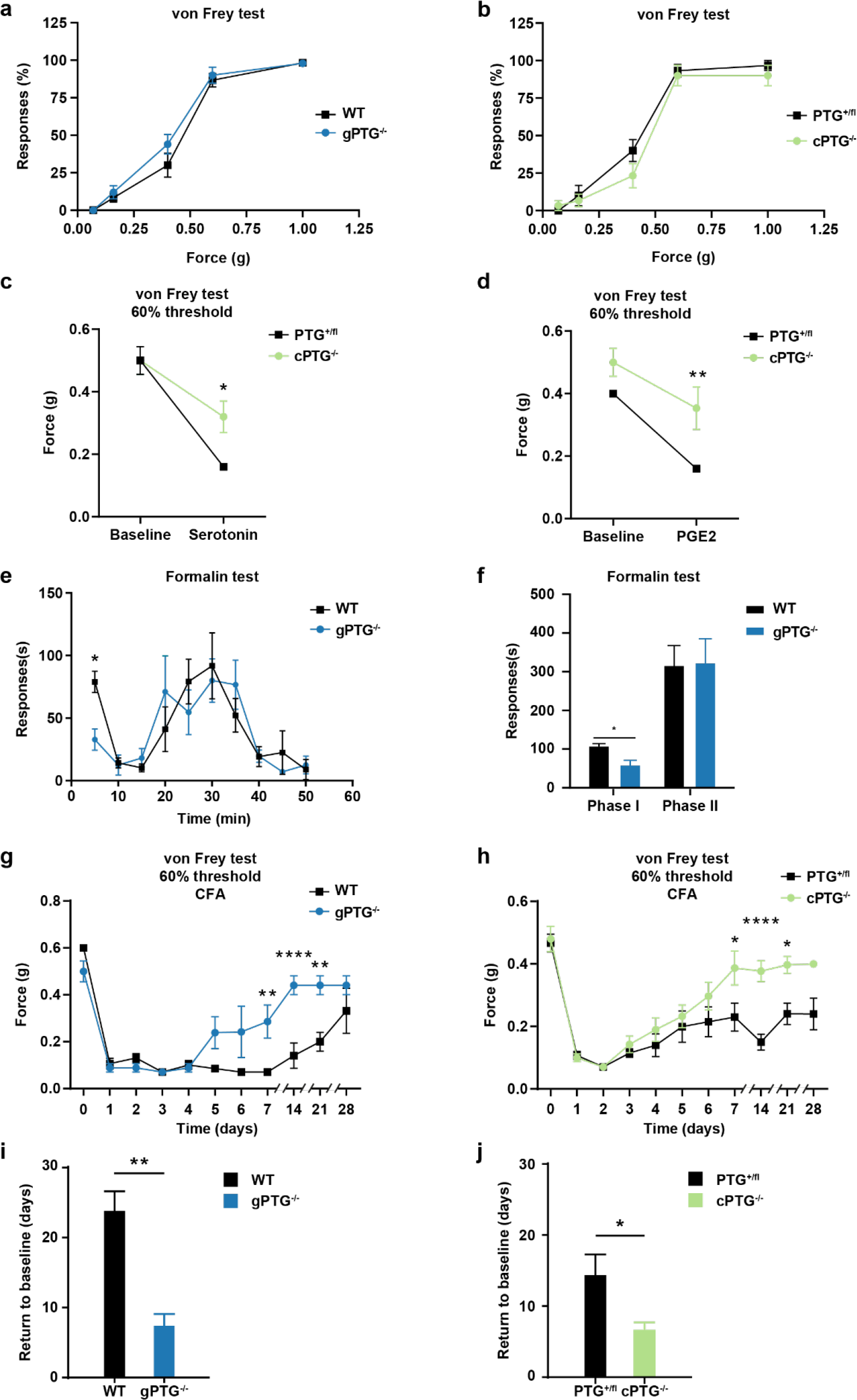
Inflammatory nociceptive sensitization and pain maintenance is reduced in PTG^(-/-)^ mice. **a,b**, Baseline mechanical sensitivity for WT and gPTG^(-/-)^ mice (**a**, N=6) or PTG^(+/fl)^ and cPTG^(-/-)^ mice (**b**, N=12). **c**,**d**, Mechanical threshold required to elicit a response in at least 60% of trials in PTG^(+/fl)^ and cPTG^(-/-)^ mice that are either not treated (baseline) or 15 minutes after intraplantar serotonin (**c**, N=6) or PGE2 (**d**, N=6) injection into the hindpaw. **e,f,** Time course of formalin-induced nocifensive responses scored in 5 min bins (**e**) or separated in Phase I (0-10min) and Phase II (10-50min) (**f**) for WT and gPTG^(-/-)^ mice (N=5). **g**,**h**, 60% mechanical threshold measured before and at different time points after CFA stimulation comparing WT (N=6) and gPTG^(-/-)^ mice (**g**, N=5) and PTG^(+/fl)^ and cPTG^(-/-)^ mice (**h**, N=12). **i**,**j**, Number of days required to recover from a sensitized state back to baseline mechanical threshold (cutoff: one standard deviation of the average baseline value) after intraplantar CFA injection of WT and gPTG^(-/-)^ mice (**i**) and PTG^(+/fl)^ and cPTG^(-/-)^ mice (**j**), data corresponds to traces shown in **g** and **h**. (A-H) Two-way ANOVA with Bonferroni post hoc test; (**i**-**j**) Unpaired two-tailed t-test. Data represent mean ± s.e.m. *p < 0.05, **p < 0.01, ***p < 0.005, ****p < 0.001. See also Extended Data Fig. 4

Collectively, these data suggest that acute spinal glycogen levels and their utilization only have a small effect on acute noxious signal processing and pain-related rodent behaviors. For the most part, short-term acute pain signaling, such as that initiated by capsaicin, appeared not to depend on glycogen turnover and was not altered in the absence of the *Ptg* gene.

The small acute pain-related phenotype in PTG^(-/-)^ animals is not entirely surprising given the observed delay to PTG induction and glycogen build-up subsequent to an inflammatory pain stimulus (**Figs. 1 and 2**) compared to duration of short-term pain sensitization.

We therefore wondered whether pathological long-term inflammatory sensitization and behavior would be altered in PTG^(-/-)^ mice. We found intraplantar CFA injections to generate strong induction in spinal glycogen accumulation that lasted up to 6 days (**Extended Data Fig. 2e**) and that sensitized the animals by lowering pain thresholds for up to 4 weeks (**Fig. 4g,h and Extended Data Fig. 4e,f**), similar to what has been shown in previous studies^24, 25^. As expected, both gPTG^(-//-)^ and cPTG^(-/-)^ animals became sensitized to mechanical and thermal stimuli similar to the wildtype controls. Strikingly, however, gPTG^(-/-)^ and cPTGA^(-/-)^ animals recovered faster from the sensitized state, reaching baseline pain thresholds already after 7±2 days compared to wildtype mice that stayed sensitized for up to 24±3 days (**Fig. 4g-j and Extended Data Fig. 4e,f**). In both PTG knock-out models, this accelerated recovery was observed for both mechanical- and heat-pain hypersensitivity, with a more pronounced beneficial effect on mechanical hypersensitivity. In summary our data demonstrate that, while short-lasting sensitization is largely unaffected in PTG^(-/-)^ mice, recovery from strong and long-lasting inflammatory pain is accelerated when astrocytic glycogen induction is blunted.

### Noxious stimuli-induced spinal glycolytic capacity and lactate production are reduced in the absence of PTG

Neuronal transmission, signal processing and plasticity require energetic metabolic support. Neurons receive energetic metabolites such as glucose via the bloodstream. However, upon increased demand, for example in the context of plasticity-triggering synaptic activity, it has been shown that neurons may depend on additional metabolic fuel from astrocytes^31–38^. We therefore wondered whether inflammatory signal processing leading to pain may modulate spinal energy consumption and, more importantly, whether astrocytic glycogen stores are a relevant fuel source. In order to measure energy consumption and respiration in the spinal network, we utilized the so-called Seahorse assay using mouse spinal cord slice preparations. This assay enables measurements of the rates of glycolysis (assessed by the <u>e</u>xtra<u>c</u>ellular <u>a</u>cidification rate ECAR) and mitochondrial respiration (assessed by the <u>o</u>xygen <u>c</u>onsumption rate OCR)^39^. To the best of our knowledge, this assay has not been described for spinal cord tissue before. Therefore, we first established spinal slice preparations and optimized measurement conditions (see methods section for details), based on a protocol established for brain slices^40^.

Next, we compared basic respiratory parameters in gPTG^(-/-)^ animals and wildtype controls. We did not find any significant differences in spinal cord tissue from these two groups of mice (**Extended Data Fig. 5a-c**). This is in agreement with results showing that astrocytic PTG and glycogen appears not to be required for basic spinal cord function because basal mechanical and thermal sensitivity are normal in PTG^(-/-)^ animals (**Fig. 4a,b and Extended Data Fig. 4a,b**).

**Fig. 5:**
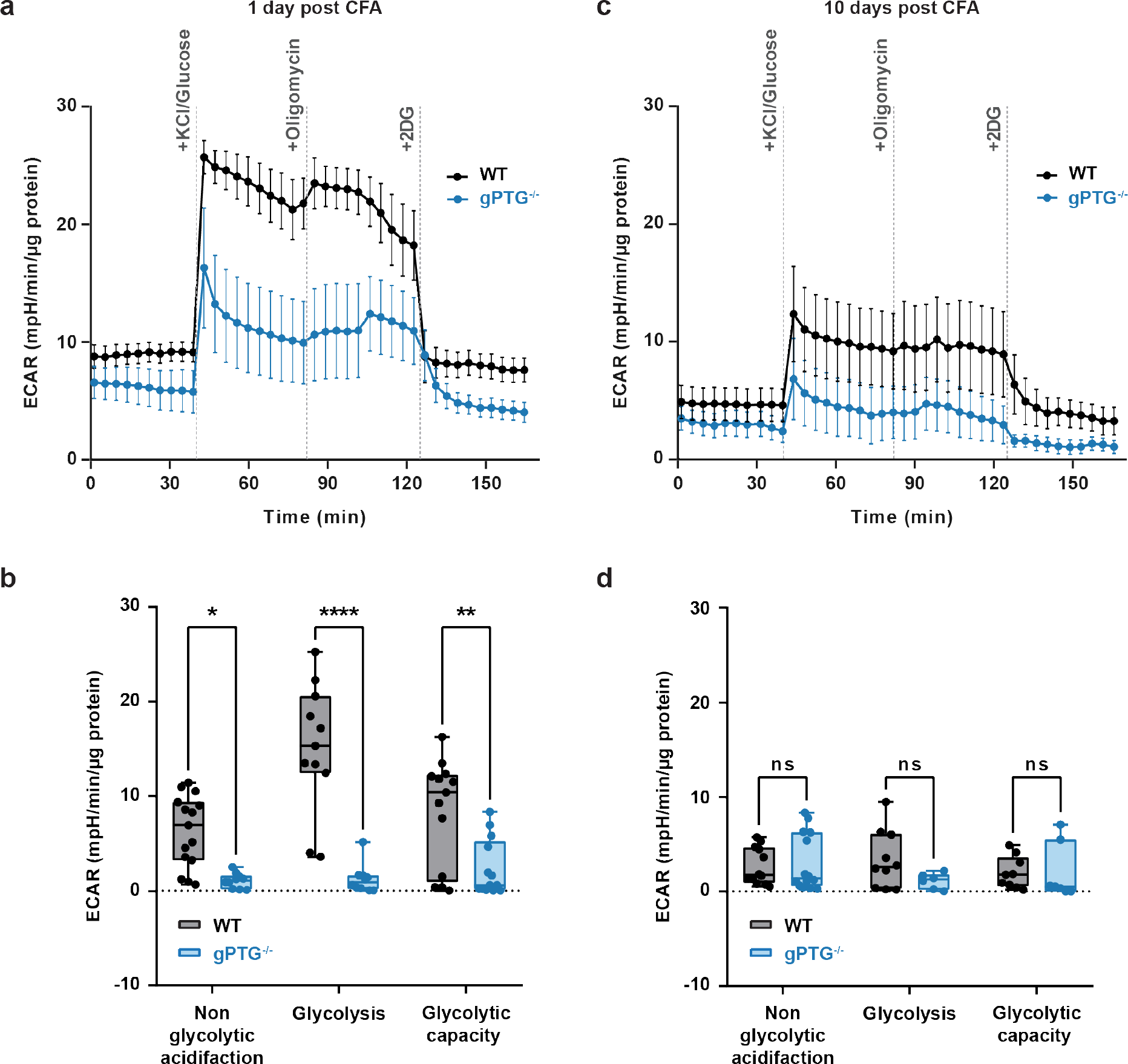
Inflammatory pain-induced glycolytic capacity in spinal cord circuits is blunted in the absence of PTG. **a,c**, Representative curves of Seahorse glycolytic rate assay during neuron stimulation induced by 25mM KCl in spinal dorsal horn slices collected 1 day (**a**) and 10 days post CFA-stimulation (**c**) of WT (N=7 samples/3 mice for 1d, N=10/3 for 10d), and gPTG^(-/-)^ (N=5/3 for 1d and N=10/3 for 10d). **b**,**d**, Comparison of non-glycolytic acidification, glycolysis and glycolytic capacity at 1 day (**b**) and 10 days (**d**) post CFA stimulation of WT mice (N=12/3 for 1d, N=7/3 for 10d), and gPTG^(-/-)^ mice (N=10/3 for 1d and N=10/3 for 10d). Two-way ANOVA with Sidak’s post hoc test. Data represent mean ± s.e.m. *p < 0.05, **p < 0.01, ***p < 0.005, ****p < 0.001. See also Extended Data Fig. 5.

We next wondered whether mimicking painful stimulation by increasing neuronal activity (action potential firing), either by supplying the TRPV1 agonist capsaicin or by perfusing spinal slices with high potassium chloride (KCl, 25mM) would trigger increased energy consumption. Indeed, both paradigms resulted in increased glycolytic activity in spinal slices obtained from wildtype mice, with KCl provoking a more robust response (**Extended Data Fig. 5d,e**). Of note, capsaicin and KCl stimulation did not result in any measurable alteration of mitochondrial oxygen consumption when tested on spinal slices of wildtype animals (data not shown).

Since we found CFA-induced inflammatory pain to trigger strong glycogen dynamics that were completely blunted in PTG^(-/-)^ mice, and because PTG^(-/-)^ mice recovered faster than wildtypes from CFA-evoked long-lasting pain hypersensitivity, we reasoned that neuronal activation on the background of CFA-induced pain would potentially reveal altered energy consumption in spinal slices of PTG^(-/-)^ mice.

We therefore subjected spinal cord slices of CFA-treated wildtype and PTG^(-/-)^ mice to glycolytic Seahorse measurements 1 day and 10 days after intraplantar CFA injection. Interestingly, the glycolytic response upon neuronal (KCl) activation was reduced in PTG^(-/-)^ spinal tissue slices obtained from animals 1 day after CFA treatment when compared to those obtained from wildtype controls (**Fig. 5a,b**). These results suggest that glycolytic capacity in the dorsal spinal cord during inflammatory pain requires elevated astrocytic glycogen stores. On the other hand, 10 days after the induction of CFA-triggered inflammatory pain hypersensitivity, glycolytic capacity was reduced for both groups of mice and appeared not significantly different between the two genotypes (**Fig. 5c,d**). These differences in glycolytic capacity correlated with differences in glycogen levels that we found to be robustly increased in wildtype (but not in PTG^(-/-)^) mice 1 day after CFA injection and that had returned back to baseline levels between 7 and 10 days after CFA injection (**Fig. 3b**).

Because spinal glycolytic capacity was reduced in PTG^(-/-)^ animals, we wondered whether levels of lactate, a downstream product of glycolysis that has been implicated in astrocyte-neuron metabolic coupling ^1^, would also be changed. Indeed, spinal lactate levels were reduced in gPTG^(-//-)^ mice (**Extended Data Fig. 5f**). Moreover, we found spinal lactate levels to be dynamically altered in spinal cord tissue of wildtype mice upon pain stimulation (**Extended Data Fig. 5g**).

Taken together, these data support the hypothesis that spinal astrocytic glycogen dynamics evoked by robust noxious stimulation and signal processing promote astrocytes’ glycolytic capacity and lactate production, thereby prolonging pain maintenance.

### Astrocytic glycogen fuels maladaptive neuronal plasticity in the spinal cord triggered by persistent noxious stimuli

How could a deficit in the energy landscape of the spinal astrocyte–neuron network be beneficial to the recovery of long-lasting pain? Persistent noxious signaling, arising for example as a consequence of tissue damage, cancer or other forms of disease, can lead to structural and functional plasticity in the spinal network, resulting in increased sensitivity and activity of spinal pain circuits and culminating in increased sensitivity to peripheral stimuli^41–44^. We hypothesized that, similar to plasticity mechanisms found in the brain^38, 45^, induction and maintenance of maladaptive pain-induced spinal plasticity requires metabolic energy that—at least in part—comes from astrocytes. To assess spinal plasticity, which is associated with heightened synaptic activity and a presumably increased propensity of spinal projection neurons to fire action potentials (APs)^46^, we first measured changes in neuronal excitability in acute spinal slice preparations of wildtype mice to assess whether this intrinsic neuronal property is enhanced in our CFA model of inflammatory pain and could serve as a measure of spinal plasticity.

To assess intrinsic neuronal excitability, we first measured the input resistance (Rin) and the rheobase of randomly sampled lamina 1 (L1) spinal neurons. The first parameter, Rin, was not changed in neurons recorded from animals 10 days after CFA injection compared to naïve control animals, suggesting that basic electrical plasma membrane properties of the neurons, such as the density of passive, non-voltage-dependent “leak” channels, did not change as a consequence of CFA treatment (**Fig. 6a**). Based on previous work, we expected the excitability of wildtype L1 neurons to increase in the wake of inflammatory pain^46^. Indeed, we found that the minimum (rheobase) current necessary to elicit an AP during a 500 ms current injection step was significantly lower in CFA-treated wildtype neurons compared to the naïve wildtype group (**Fig. 6b**), establishing this cell intrinsic electrophysiological parameter as a bona fide inflammatory pain-induced spinal plasticity indicator.

**Fig. 6:**
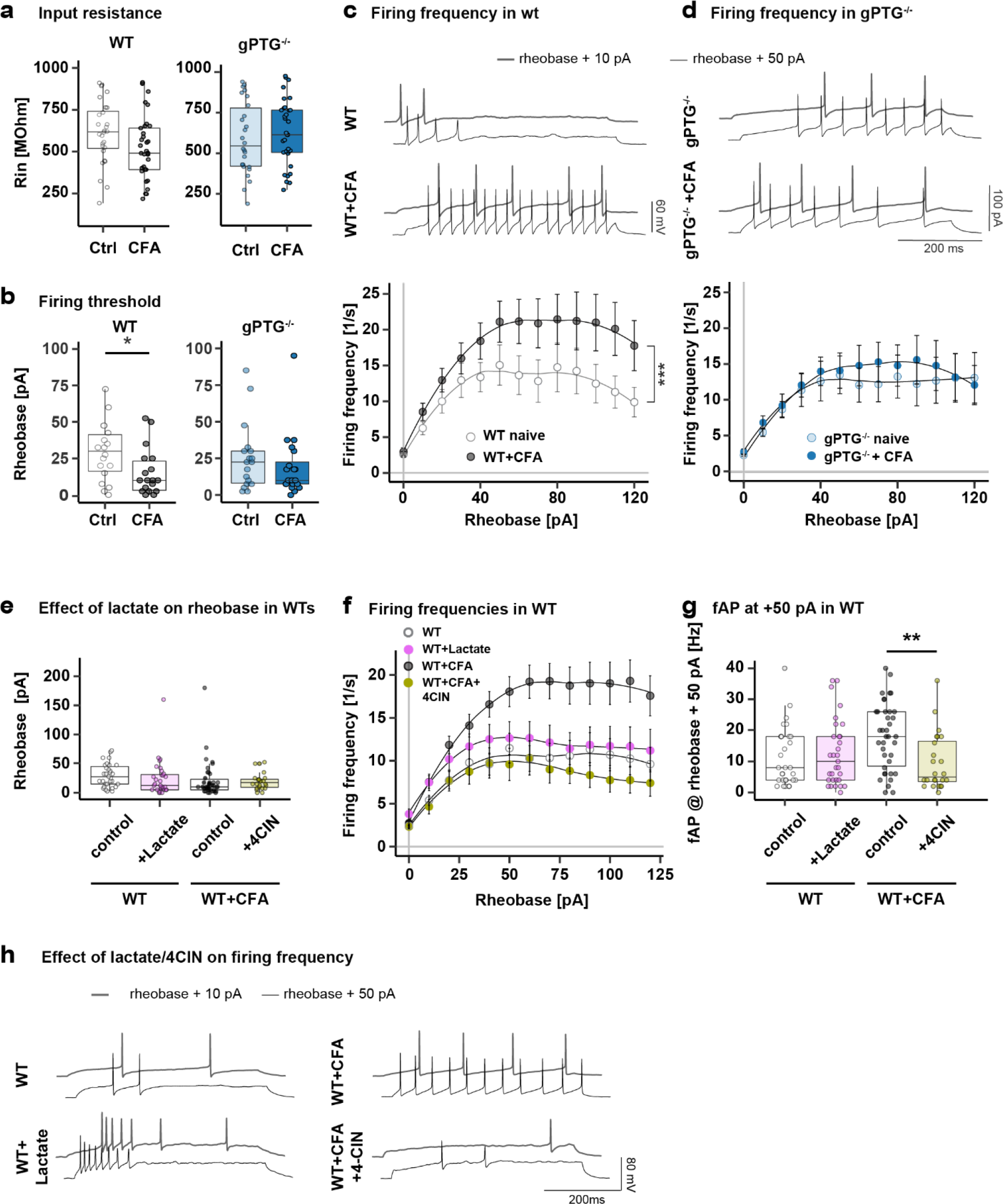
CFA-induced spinal neuronal plasticity is blunted in gPTG^(-/-)^ mice. **a**, Comparison of membrane input resistance (Rin) of randomly recorded L1 spinal neurons derived from naïve and CFA-injected (10 days post-injection) WT and gPTG^(-/-)^ KO mice. n=28 cells/N=4 animals (WT), n=33/4 (WT+CFA), n=26/4 (KO) and n=33/4 (KO+CFA). **b**, Comparison of the rheobase in L1 neurons from naïve and CFA-treated WT and gPTG^(-/-)^mice. n=18 cells/N=4 animals (WT), n=18/4 (WT+CFA), N=18/4 (gPTG^(-/-)^) and n=20/4 (KO+CFA). Unpaired two-tailed t-test, *p = 0.0283 for the WT condition. **c**, Top: Example traces of firing patterns in L1 neurons derived from WT naïve and CFA-treated mice. Bottom: Comparison of the firing frequencies across a range of 500ms-long current injections (from 0pA to 120pA above rheobase) in WT and WT+CFA group. n=18 cells/N=4 animals (WT) and n=20/4 (WT+CFA). Two-way ANOVA (effect of CFA treatment), ***p < 0.001. **d**, Top: Example traces of firing patterns in L1 neurons derived from wildtype naïve and CFA-treated mice. Bottom: Comparison of the firing frequencies across a range of 500ms-long current injections (from 0pA to 120pA above rheobase) in gPTG^(-/-)^ and gPTG^(-/-)^+CFA group. n=18 cells/N=4 animals (gPTG^(-/-)^) and n=20/4 (gPTG^(-/-)^ +CFA). **e**, Rheobase comparison of WT/WT+CFA groups recorded after the addition of 4-CIN (300μM) or lactate (15mM) into recording solution (aCSF). n=31 cells/N=4 animals (WT), n=31/3 (WT + Lactate), n=38/4 (WT+CFA) and n=24/3 (WT+CFA+4-CIN). **f**, Comparison of the firing frequencies in response to 500ms current injections (from 0pA to 120pA above t rheobase) in naïve/CFA WT groups in the presence of L-lactate or 4-CIN. **g**, Comparison of firing frequencies at 50pA above rheobase (based on **f**); n=30 cells/N=4 animals (WT), n=31/3 (WT+Lactate), n=38/4 (WT+CFA) and n=24/3 (WT+CFA+4-CIN). One-way ANOVA, p<0.0001; Sidak’s post hoc test, **p=0.003 (WT+CFA:WT+CFA+4-CIN). **h**, Example traces of firing patterns of WT/WT+CFA neurons recorded in the presence of 4-CIN or lactate. Data are shown as mean ± s.e.m. ∗p < 0.05, ∗∗p < 0.01, ∗∗∗p < 0.001. See also Extended Data Fig. 6.

Strikingly, CFA treatment did not affect the rheobase in PTG^(-/-)^ neurons 10 days after intraplantar injection (**Fig. 6b**), suggesting that blunted glycogen dynamics prevented hyperexcitability. We further analyzed the neuronal excitability of L1 neurons by injecting a range of depolarizing currents above the threshold (rheobase) current to evoke trains of APs. Again, we found that CFA treatment resulted in enhanced AP firing to current injections in wildtype neurons (**Fig. 6c**) but not in neurons derived from gPTG^(-/-)^ animals (**Fig. 6d**).

Because our metabolic analysis suggested that lactate, a downstream metabolite of glycogenolysis/glycolysis, might play a role in mediating hyperexcitability, we next set out to test whether the firing properties of L1 spinal neurons can be modulated by either application of α- cyano-4-hydroxycinnamate (4-CIN, 300µM), an inhibitor of monocarboxylate transporters involved in shuttling lactate between astrocytes and neurons^47^ or by supplementation of lactate in the perfusion fluid. 4-CIN did not affect rheobase in L1 neurons from either wildtype naïve or wildtype mice 10 days post CFA injection (**Fig. 6e**). Interestingly, however, the drug robustly reduced CFA-induced hyperexcitability in wildtype L1 neurons back to baseline levels when neurons were stimulated to fire trains of action potentials (**Fig. 6f-h**). Conversely, at concentrations shown to enhance firing activity of cortical neurons (15 mM)^45^, lactate supplementation did not have any stimulatory effect on the firing properties of naïve wildtype neurons, suggesting that lactate alone is not sufficient to transform L1 neurons into an hyperexcitable state.

Since we found L1 neurons from gPTG^(-/-)^ animals to be compromised in increasing their induced AP firing rates following CFA treatment, we assessed whether adding lactate to gPTG^(-/-)^ mouse-derived spinal slices would be sufficient to increase their excitability. However, again lactate alone did not affect rheobase in gPTG^(-/-)^ slices, neither from naïve nor CFA-treated animals (**Extended Data Fig. 6a**). Similarly, lactate supplementation did not majorly modulate firing frequencies in L1 neurons in response to current injections above rheobase (**Extended Data Fig. 6b,c**).

Together, these results show that astrocytic glycogen dynamics is an important factor driving hyperexcitability of dorsal spinal L1 neurons in the context of inflammatory pain. While lactate transfer via monocarboxylate transporters to neurons appears to play some role in regulating hyperexcitability, acutely supplementing lactate 10 days after spinal plasticity has been triggered by CFA treatment is not sufficient to promote hyperexcitability in PTG^(-/-)^ mice.

Our data suggest that PTG and glycogen dynamics are important contributors to maladaptive spinal plasticity and the maintenance of long-lasting pain hypersensitivity. Since beneficial forms of plasticity in higher brain centers ––associated with learning and memory–– have also been shown to depend on astrocytic metabolism, we wondered whether PTG^(-/-)^ mice are compromised in cognitive memory formation. We therefore assessed the ability of the mice to perform in the Morris Water Maze test, a classic paradigm to assess spatial learning and memory in rodents^48^. We found that PTG^(-/-)^ mice performed similar to wildtype control mice in this assay (**Extended Data Fig. 7**), suggesting that the metabolic PTG pathway does not majorly contribute to this learning and memory paradigm.

## Discussion

Brain astrocytes have been shown to provide metabolites and metabolic energy for neighboring neurons in situations when additional energy is required, such as forms of long-term plasticity and memory formation^31–38^.

Plasticity of spinal neurons, such as long-term potentiation and enhanced excitability of lamina 1 neurons of the pain pathway, contributes to persistent forms of pain^42, 44, 46, 49^. While the induction and acute phase of spinal plasticity has been researched extensively and several molecular mechanisms have been identified, mechanisms prolonging this state of plasticity and thus enabling transition to persistent pain are yet to be fully understood.

Astrocytes in the spinal cord have been shown to mediate long-term nociceptive sensitization in particular in models of inflammatory pain^6, 8^. Together with microglia, astrocytes have also been described to contribute to pain-induced, glia-mediated forms of spinal plasticity^50^. Importantly, while glial contributions have been identified in terms of molecular signaling and release of pro-inflammatory mediators, they have not been tested in terms of driving metabolic plasticity relevant to pain.

We show here that spinal astrocytes, triggered by painful stimuli, adjust their energy metabolism to sustain and maintain long-lasting inflammatory pain states.

Previous work has provided evidence that metabolic inhibitors blocking the tricarboxylic acid (TCA) cycle, such as fluorocitrate and fluoroacetate, are effective at inhibiting pain in rodent models. Intriguingly, these metabolic blockers are preferentially taken up by astrocytes, suggesting that the metabolic perturbation of astrocytes can have beneficial effects and alleviate persistent forms of pain^6, 51^. However, these drugs are only partially selective and, eventually, result in astrocyte decline, complicating the interpretation of these results.

Our study uncovers a genetically-defined pathway that, unexpectedly, drives glycogen dynamics in spinal astrocytes subsequent to inflammatory pain stimulation. Noxious stimulation-induced acute glycogen mobilization, subsequent PTG induction and glycogen build-up, follow a long-term trajectory that scales with the magnitude/severity and duration of the inflammatory pain stimulus and that, according to our data, is less relevant for the early stages of pain states but more important for the maintenance of long-term inflammatory pain. We show that blunting PTG-driven spinal glycogen dynamics reduces nociceptive neuronal plasticity in the spinal cord and, presumably as a consequence, shortens the maintenance phase of inflammatory pain states, as assessed in behavioral experiments.

Our energy metabolic measurements and electrophysiological recordings conjointly emphasize robust neuronal-astrocyte coupling, whereby pain-induced neuronal activity (directly or indirectly) triggers astrocytic glycogen dynamics, thereby promoting increased glycolytic capacity to reinforce pain maintenance.

We can only speculate about the nature of such a signal that triggers the metabolic changes in astrocytes. It is possible that dorsal spinal astrocytes respond directly to neurotransmitters released by primary afferent fibers, such as glutamate and/or inflammatory mediators^6^. Interestingly, it has been shown recently that afferent pain signals acutely activate spinal astrocytes indirectly via the release of noradrenaline (NA) from descending fibers originating in the Locus Coeruleus^8^. NA, but also insulin, glutamate and other soluble signaling molecules can stimulate an increase in glycogen content in cultured astrocytes^52–57^, a property that *in vitro* has been linked to PTG induction^23^. It is therefore conceivable that neuronal activity of primary afferent nociceptive neurons promotes astrocytic glycogen build-up by a soluble signaling molecule released in the spinal cord.

But how can glycogen build-up result in enhanced glycolytic capacity in the spinal cord as we observed in extracellular acidification rate (ECAR) and oxygen consumption rate (OCR) measurements? A potential explanation is offered by a recent study that analyzed PTG^-/-^ adipocytes^58^. Similar to astrocytes, in adipocytes NA also promotes PTG-dependent glycogen build-up. However, it was found that not the build-up per se, but rather the build-up coupled with glycogen turnover ––glycogen dynamics–– is required for cellular function, an idea that previously had been hinted at by studies that detected glycogen “over-accumulation” in adipose tissue^59, 60^. We hypothesize that a similar phenomenon also prevails in spinal astrocytes in the course of pain signal processing: It is increased glycogen dynamics, rather than solely glycogen accumulation, that is relevant for the cellular and behavioral phenotypes we observe. This may explain why we find glycolytic capacity to be reduced in PTG^-/-^ spinal cord tissue: the lack of noxious stimulation-induced glycogen build-up in the absence of PTG results in reduced substrate availability for glycolysis (glucose generated via glycogenolysis) and thereby prevents an increase in glycolytic capacity compared to the wildtype dorsal spinal cord network.

In such a scenario, spinal glycogen dynamics would act as a metabolic buffer and noxious stimuli- boosted glycolytic capacity in the spinal cord of wildtype mice would provide metabolites that may signal and/or calorically-fuel spinal plasticity.

Among the different metabolites that are shuttled between astrocytes and neurons, lactate has received considerable attention. Since the 1990’s it has been postulated that astrocytic glycogen may constitute a source for lactate (rather than for glucose) for neighboring neurons^32^. Lactate can enhance excitability of some neurons acutely^61^, possibly also independently of its caloric value^62^ and the metabolite was found to foster neuronal plasticity and memory formation in higher brain centers^38^.

Moreover, enhanced long-term excitability in wildtype spinal L1 neurons was found to be reduced by inhibiting lactate transporters present in neurons and astrocytes^10^.

Because our electrophysiological data also pointed to a reduction of plasticity and diminished noxious stimulation-induced excitability in the absence of glycogen dynamics, we considered lactate as one potential metabolite induced to shuttle from astrocytes to spinal neurons as a consequence of long-term pain stimulation. Additionally, our metabolomics analysis suggested that spinal lactate levels are indeed dynamically altered during pain stimulation and are reduced in PTG^(-/-)^ mice. Indeed, blocking monocarboxylate (lactate) transporters reduced inflammatory pain induced hyperexcitability in spinal L1 neurons. However, lactate supplementation to PTG^-/-^ spinal slices obtained from mice stimulated with CFA was not sufficient to promote excitability.

These results suggest that lactate, mobilized in the process of glycogen dynamics and possibly shuttled from astrocytes to neurons during the late, maintenance phase of inflammatory pain, may constitute a metabolite required for enhanced excitability in L1 neurons. But lactate is not sufficient and requires other plastic changes to occur in the wake of an inflammatory insult in order to act either as a signaling molecule or as metabolite with caloric value to booster hypersensitivity in spinal L1 neurons.

Future work, requiring cell-type selective metabolic flux measurements and refined metabolic *ex vivo* and *in vivo* perturbations, combined with neuro-glia activity and plasticity measurements, will help to shed light onto this clinically highly relevant phenomenon, which may extend to other CNS circuits involved in pain signal processing.

Interestingly, protein phosphatase 1 (PP1), the signaling molecule regulated by PTG to control glycogen content, has previously been implicated in neuronal plasticity in higher brain centers to foster learning and memory^63–66^. We therefore wondered whether PTG^(-/-)^ mice would have deficits in learning and memory. However, we did not find any gross abnormalities when subjecting PTG- deficient animals to the Morris Water Maze test, suggesting that other metabolic mechanisms are at play to fuel these types of plasticity. Given that we find glycogen dynamics in spinal astrocytes to scale with the magnitude of the pain stimulus, we speculate that more subtle sensory stimuli required to foster learning and memory in higher brain centers ––and following different kinetics compared to long-lasting noxious stimuli–– likely correlate with subtle changes in the energetic landscape that are independent (or less dependent) on the transcriptional induction of PTG.

We believe these findings will fuel further research to uncover a potential new class of analgesics, specifically designed to interfere with neuron-glia metabolic coupling and thereby blocking the maladaptive neuronal plasticity associated with the maintenance of pain states.

## Methods

### Mice

All animal care and experimental procedures were approved by the local council (Regierungspräsidium Karlsruhe, Germany) under protocol numbers 35-9185.81/G-168/15, 35- 9185.81/G-201/16, 35-9185.81/G-295/21, 35-9185.81/G-173/21. Mice were housed under standard conditions with a 12h light/dark cycle and ad libitum access to food and water. All genetically modified mice in this study were on the C57BL/6N background. All mice were bred at the animal facility of the Heidelberg University (Interfakultäre Biomedizinische Forschungseinrichtung, IBF) or purchased from Janvier Labs. Both male and female mice were used for the experiments. The animals were randomly assigned for experiments and the experimenter was aware of the animal genotype when conducting experiments except for behavior tests. Between 7- and 16-weeks old mice were used, except for Seahorse experiments where mice were 3-5 weeks old. Prior to experiments, mice were given 24-72h to habituate in the laboratory.

### Mouse lines used

All Wild type animals used in this study correspond to the C57BL/6N mouse line.

cPTG^(-/-)^ and gPTG^(-/-)^ mouse lines were made as part this study using standard transgenic methods (see Extended Data Fig. 3a for details). In brief, after generating PTG^(+/fl)^ mice, these were crossed with a deleter-Cre strain provided by Reinhard Fässler from Max-Planck-Institut für Biochemie (Takahashi et al., 2007) to eliminate the loxP flanked region, generating PTG^(+/-)^ animals. gPTG^(-/-)^ where then generated by inbreeding of PTG^(+/-)^.

Astrocyte-specific conditional PTG KO animals, cPTG^(-/-)^, were obtained by crossing two lines: First, a homozygous floxed PTG line (PTG^(fl/fl)^) was generated by inbreeding of PTG^(+/fl)^ giving PTG^(fl/fl)^ mice as a result.

Second, gPTG^(-/-)^ mice were crossed with an astrocyte specific Tamoxifen-dependent line (Aldh1L1-Cre^ERT2^) obtained from the laboratory of Bal Khakh at University of California, Los Angeles (Srinivasan et al., 2016) generating Aldh1L1-Cre^ERT2^-PTG^(+/-)^ mice. Finally, this line was crossed with PTG^(fl/fl)^, producing Aldh1L1-Cre^ERT2^-PTG^(fl/-)^ mice.

cPTG^(-/-)^ were finally generated by injecting i.p 200 µL of a solution containing 10mg/mL Tamoxifen (Sigma, T56485) dissolved in a mixture of 5% ethanol and 95% sunflower oil (Sigma, 88921). Animals were injected once a day for 5 consecutive days. Littermates PTG^(+/fl)^ that did not express Aldh1L1-Cre^ERT2^ were injected with tamoxifen at the same time and used as control group for our experiments. Experiments were conducted only 3 weeks after the last injection.

### Pain models

<u>Formalin pain model:</u> 36.5% formaldehyde solution (Sigma, F8775) was diluted with 0.9% NaCl sterile saline (B.Braun, 190/150936) to a 5% solution shortly before injection. Formalin-induced pain was released by intraplantar injection of 15 µL of the 5% formaldehyde solution into the hind paw of the mouse, under shallow isoflurane anesthesia.

<u>Capsaicin pain model:</u> 0.6% capsaicin/DMSO solution (obtained from the laboratory of Rohini Kuner) was diluted with sterile saline to a 0.06% suspension shortly before injection. Capsaicin-induced pain was released by intraplantar injection of 20 µL of the 0.06% capsaicin solution into the hind paw of the mouse, under shallow isoflurane anesthesia.

<u>Serotonin pain model:</u> 0.5% Serotonin/0.9% NaCl solution. Serotonin-induced pain was released by intraplantar injection of 20 µL of the Serotonin solution into the hind paw of the mouse, under shallow isoflurane anesthesia.

<u>PGE2 pain model:</u> 2.5 ng/µL 0.9% NaCl solution. PGE2-induced pain was released by intraplantar injection of 20 µL of the PGE2 solution into the hind paw of the mouse, under shallow isoflurane anesthesia.

<u>CFA pain model:</u> 20 µL of Complete Freund’s Adjuvant (Sigma, F5881) was injected into the hind paw while the mouse was kept under shallow anesthesia by isoflurane, to induce inflammatory pain.

<u>SNI pain model:</u> The SNI surgery was performed by severing two of the three branches of the sciatic nerve (the tibial nerve and the common peroneal nerve). The mice were kept under isoflurane anesthesia during surgery an let to recover for at least one week before subjected to any measurement/experiments.

### Immunofluorescence Staining

<u>Sample preparation:</u> At different time points after intraplantar injection of formalin, mice were perfused first with PBS (Sigma, D8537) and then 4% Paraformaldehyde/PBS (Sigma, 16005). After perfusion, mice were dissected to collect the lumbar spinal cord. The spinal cord piece was incubated in 20% sucrose/PBS (Sigma, S0389) overnight at 4°C with gentle shaking for cryoprotection and then embedded in OCT (Tissue-Tek® O.C.T. Compound, Sakura, #4583). The spinal cord sample was sectioned into 12 µm slices on a Leica CM3050S Research Cryostat and sections were collected on glass slides (HistoBond, #0810001). Sections were dried at 37°C and then stored at -80°C until further use.

<u>Staining procedure:</u> Immunofluorescent stainings were performed on spinal cord sections collected as described above. First, the sections were equilibrated to room temperature and then washed twice for 5 minutes (min) in PBST (0.1% Triton X-100 (Merck, #108603)/PBS). Blocking was performed with 1% goat serum in PBST for 1 hour at room temperature (RT). The sections were incubated with primary antibody (pS6 Cell Signaling 2215, 1:1000; GFAP Cell Signaling #3670, 1:500; NeuN Cell Signaling D4G40, 1:2000; IBA1 Wako, #019-19741, 1:500) in blocking buffer overnight at 4°C. Next day, the slices were washed 3 times in PBS-T for 10 min each time, before secondary antibody (Donkey α-Rabbit, Alexa Fluor 488, Invitrogen #A21206, 1:2000; Goat α-mouse, Alexa Fluor 488, Invitrogen #A11001, 1:2000 in blocking buffer) was applied to the sections and incubated for 1 h at RT. The sections were washed 3 times in PBST for 10 min each time, mounted in Immu-Mount (Thermo Scientific, #9990402) and stored at 4°C until imaging.

### Wholemount immunofluorescence staining

<u>Sample preparation</u>: Mice were sacrificed 2 h after intraplantar formalin injection and immediately perfused as previously described. After perfusion, the spinal cord was dissected out and subjected to 4h of post-fixation in 4% Paraformaldehyde/PBS at 4°C while gentle rotating. The tissue piece was then washed in PBS for 10 min and put into Dent’s Bleach (10% H_2_O_2_(Merck, #107209), 13.3% DMSO (Sigma, D2438) and 53.3% Methanol (Honeywell, #32213)) for 24h at 4°C while gentle rotating. After bleaching, the tissue piece was dehydrated using methanol, replacing it 5 times after 2 minutes each time. The sample was then fixed in Dent’s Fix (20% DMSO, 80% Methanol) for 24 h at 4°C with gentle rotation and then stored in Dent’s Fix at 4°C until further use.

<u>Staining procedure:</u> The processed tissue was rinsed 3 times with PBS (20min each) to wash off the Dent’s Fix and then incubated with primary antibody in blocking buffer (pS6 Cell Signaling 2215, 1:1000) at 4°C with gentle rotation for 5 days. After primary antibody incubation, the tissue sample was washed 6 times with PBS (30min each time) at RT. Secondary antibody (diluted in blocking solution) was (Donkey α-Rabbit, Invitrogen #A21206, 1:2000) and incubated for 2 days at RT while gentle rotating. After the incubation, the tissue was washed 6 times with PBS (30min each time) at RT.

<u>Clearing procedure:</u> After staining, the spinal cord piece was dehydrated and cleared before imaging. Dehydration was done by incubating the tissue in 50% methanol/PBS for 5 min at RT, followed by 20 min incubation in 100% methanol, changing the solution twice during the incubation. After dehydration, the tissue was cleared in BABB (1 part benzyl alcohol (Sigma, 108006) and 2 parts benzyl benzoate (Sigma, B6630)) for 5 min and stored in BABB until imaging.

### Immunoprecipitation/Ribosome capture

<u>Sample preparation:</u> For sequencing experiments, 5 mice were pooled for each sample. Three samples of each condition were prepared. Immunoprecipitation was performed based on the protocol developed by Knight et al., 2012. In brief: Mice were sacrificed 2hs after formalin injection and quickly dissected into freshly prepared, ice-cold buffer B (1xHBSS (Life Technologies, #14185-045), 4 mM NaHCO3 (AppliChem, #131638.1211), 2.5 mM HEPES (pH7.4; Roth, #9105.4), 35 mM Glucose (Merck, #08337), 100 µg/ml cycloheximide (Sigma, C7698)). The tissue was snap frozen and kept in -80°C until homogenization.

<u>Homogenization</u>: Tissue was resuspended in 1350 µL freshly prepared buffer C (10 mM HEPES (pH7.4), 150 mM KCl (Sigma, P9541), 5 mM MgCl2 (Sigma, M2670), 1x PhosSTOP phosphatase inhibitor tablet (Sigma, 4906845001), 1x cOmplete™ Protease Inhibitor Cocktail (Roche, 11697498001), 2 mM DTT (Sigma, 10197777001), 100 U/ml RNasin (Promega, N2515), 100 µg/ml cycloheximide)) and homogenized at 4°C. The homogenates were centrifuged at 2000g for 10min at 4°C and the supernatants (∼1mL) were transferred to fresh tubes on ice. 90 µl of 10% NP40 and 90 µl of freshly prepared 1,2-diheptanoyl-sn-glycero-3-phosphocholine (DHPC, Avanti Polar Lipids, #850306, 100mg/0.69ml) were added to each sample. The mix was centrifuged at 17000g for 10min at 4°C and the supernatant was transferred to a fresh tube. A small aliquot of the supernatant was taken for western blot (“Ly”) and the rest was used for immunoprecipitation.

<u>Antibody coating</u>: 3 µg of pS6 (Cell Signaling, clone 5364) antibody and 50 µl Protein A Dynabeads (Invitrogen, #1001D) were used for each immunoprecipitation. Antibody was incubated with Dynabeads in 1 ml buffer A (0.1% Triton X-100/PBS) for 10min at RT with gentle rotation. The coated beads were equilibrated in buffer C (see above) until immunoprecipitation.

<u>Immunoprecipitation:</u> Immunoprecipitation was performed by applying the supernatant to pS6 coated protein A Dynabeads and rotated at 4°C for 10 min. After immunoprecipitation, the liquid was collected for western blot (“SN”). The beads were washed 4 times in 500 µl buffer D (10 mM HEPES (pH7.4), 350 mM KCl, 5 mM MgCl_2_, 2 mM DTT, 1% NP40, 100 U/ml RNasin, and 100 µg/ml cycloheximide). During the third wash, the beads were transferred to a fresh tube and incubated at RT for 10 min. Phospho-ribosomes were eluted with 350 µl RLT Buffer (Qiagen, 79216) or the eluates were immediately processed to RNA extraction for test qPCR or deep sequencing experiments. Alternatively, to elution, for “IP” Western blot 50 µl 2x Laemmli was added to the Phospho-ribosomes.

### RNA extraction

RNA extraction was performed using RNAeasy micro kit (QIAGEN, #74004), following the manufacturer’s instruction. The elution volume was 10 µl. The RNA was reverse transcribed using SuperScript III Reverse Transcriptase (Thermo Fisher, #18080044) and Oligod(T)23VN (NEB, S1327) following the manufacturer’s instruction for SuperScript First-Strand Synthesis System (Thermo Fisher).

### RNA sequencing

RNA was extracted from IP eluates as described above. The obtained RNA was snap frozen and kept at -80°C until further use.

<u>Reverse transcription and amplification:</u> cDNA synthesis and amplification were performed following the SMART-Seq2 protocol described by Picelli et al., 2014 and optimized by EMBL Genecore. 2 µl of RNA was used as starting material and an 18 cycles amplification was performed.

<u>Quality control:</u> The amplified cDNA was subjected to concentration measurement using the Qubit dsDNA HS Assay kit (Thermo Fisher, Q32854) and a quality check using a bioanalyzer (Agilent High Sensitivity DNA Kit, #5067-4626). A small aliquot of the amplified cDNA was used for a qPCR testing while the remaining cDNA was kept at -20°C until further processing.

<u>Library preparation, barcoding and sequencing:</u> Library preparation and barcoding were performed following the standard protocol of EMBL Genecore (Nextera XT DNA Library Preparation Kit, FC-131-1096 and Nextera XT Index Kit v2 Set A, FC-131-2001), using 125 pg amplified cDNA as input. The sequencing was performed by EMBL Genecore on Illumina MiSeq platform.

### qPCR

qPCRs were performed using the FastStart Essential DNA Green Master (Roche, #06402712001) on a Roche LightCycler® 96 Instrument, following the manufacturer’s instruction.

qPCR primers used:

*Ptg*-Fw: GGTGACTCATCTTTCTGCCACA *Ptg*-Rv: CAAGACAAAATTAGGCACGAGA *Tubb3*-Fw: TGAGGCCTCCTCTCACAAGT *Tubb3*-Rv: GTCGGGCCTGAATAGGTGTC

### *In situ* hybridization

*Sample preparation:* Spinal cord were dissected from naïve animals or 2hs after stimulus (formalin, capsaicin, or CFA) injection into ice cold PBS and immediately embedded in OCT and frozen using dry ice/isopropanol. 20 µm sections were prepared using a Leica CM3050S cryostat, dried at 37°C and stored at -80 °C until further use.

<u>Probe design and synthesis:</u> *In situ* hybridization probes were designed using commercial software

OLIGO (version 7).

<u>Primers used:</u>

*Ptg*-Fw: GACGTCGACAAGAACTTTGTCTGCCTCGAGA

*Ptg*-Rv: GACGCGGCCGCTACCACAGCGTTCCATCACC

Probe sequences were amplified from spinal cord cDNA, molecularly cloned into pBlueScript plasmids and bacterially amplified. DIG-labeled *in situ* probes were subsequently synthesized. In situ hybridization was performed using standard procedures (Watakabe et al., 2007). In brief, DIG- and/or FITC-labeled cRNA probes were used for hybridization on cryosections. Hybridization was performed overnight at 65 °C. Sections were washed at 60 °C twice in 2xSSC/50% Formamide/0.1% N-lauroylsarcosine, treated with 20 µg/ml RNAse A for 15 min at 37 °C, washed twice in 2xSSC/0.1% N-lauroylsarcosine at 37 °C for 20 min and twice in 0.2xSSC/0.1% N-lauroylsarcosine at 37°C for 20 min. Sections were blocked in MABT/10% goat serum/1% Blocking reagent (Roche, cat# 11096176001). For NBT/BCIP staining sections were incubated overnight with sheep anti-DIG-AP (1:1000, Roche 11093274910). After washing staining was performed using NBT/BCIP in NTMT until satisfactory intensity. Double fluorescent in situs were stained by two consecutive rounds of TSA amplification with intermediate peroxidase inactivation. Sections were incubated with sheep anti-FITC-POD (1:2000, Roche cat# 18 11426346910) or sheep anti DIG-POD (1:1000, Roche cat# 11207733910). Subsequently sections were incubated with Streptavidin-Cy2 and DAPI in blocking solution, washed and mounted in Immu-Mount (Shandon).

Immunofluorescent in situ hybridization double staining: single color fluorescent *in situ* hybridization was performed following the same protocol as above: after single color fluorescent *in situ* hybridization was completed, the sections were subjected to antibody immunofluorescent staining (pS6 Cell Signaling 2215, 1:1000; GFAP Cell Signaling #3670, 1:500; NeuN Cell Signaling D4G40, 1:2000; IBA1 Wako, #019-19741, 1:500) using the protocol cited above.

### RNAscope®

Sample preparation was followed similarly to the *in situ* hybridization protocol, with the modification that sections were prepared with a thickness of 14 µm. RNAscope^®^ in situ hybridization assay was performed according to manufacturer’s instructions (Advance Cell Diagnostics (ACD), Hayward, CA, USA). A custom PTG probe (Cat No. 838311) was designed and produced by ACD.

### Glycogen assay

<u>Sample preparation:</u> Animals were sacrificed under Isoflurane anesthesia; Lumbar spinal cord dorsal horn was dissected and immediately snap frozen. The samples were kept at -80°C until further processing.

For Brain sections, animal was sacrificed under Isoflurane, the brain was quickly released from the skull and immediately frozen with dry Ice. Insula, Amygdala, Prefrontal cortex and S1 Hind Limbs sections were collected following measurements from the Allen Brain Atlas and samples were snap frozen and kept at -80°C until further processing.

<u>Glycogen measurement:</u> Tissue was sonicated in 120 µl ddH2O and incubated on a temperature shaker at 99°C for 10 min at 350 rpm to inactivate enzymes. After heat inactivation, the homogenates were centrifuged at 18000 g for 10 min at 4°C. The supernatants (100-110 µl) were collected in fresh tubes. Collected supernatants (80µl) was mixed with 34.3 µl hydrolysis buffer from a Glycogen Colorimetric/fluorometric Assay Kit (Biovision™, #K646-100), centrifugated at Room temperature for 10 min at 18000 g and 100 µL of the supernatant was divided in to wells of a 96 well plate and incubated with (sample) and without (negative control) 1 µl hydrolysis enzyme, respectively. The remaining steps were performed following the manufacturer’s instruction. A glycogen standard curve was prepared for each experiment to counter the kit-to-kit variances. 10 µl of the remaining supernatants were used for protein measurement (Pierce™ BCA Protein Assay Kit, #23225) following manufacturer’s instructions. After glycogen and protein measurements, the glycogen level of each sample was calculated by extracting the respective negative control and normalized to the protein content.

### Lactate assay

10 µl of the supernatant obtained after the first centrifugation at 18000g from the Glycogen assay where combined with 90 µl of Lactate assay Buffer (Abcam^®^ L- Lactate Assay kit, #ab65331) and separated in two wells of a 96 well plate for sample and sample background control. Lactate standard and remaining steps were performed following the manufacturer’s instruction.

### Metabolomic Analysis

Spinal cord dorsal horn was dissected as described above. Samples were immediately snap frozen and 3 samples were pooled per tube. The samples were kept at -80°C until being shipped in dry ice and processed by Metabolon^®^, USA.

### Behavioral tests

For behavior studies, littermates were used, and the experimenter was blinded to the genotype. Mice were brought into the behavioral room half an hour before beginning with the behavioral tests.

### Mechanical stimulation of mouse hindpaws (von Frey test)

Von Frey filaments (North Coast Medical, Gilroy, CA, USA) with increasing bending forces of 0.07, 0.16, 0.4, 0.6, 1, and 1.4 g were consecutively applied to the plantar surface center of both hind paws. Mice were kept in standard plastic modular enclosures on top of a perforated metal platform (Ugo Basile, Varese, Italy), enabling the application of von Frey filaments from below. Mice were acclimatized on three consecutive days for 1.5 h to the von Frey grid and 30min before starting the measurements on an experimental day. Each filament was tested five times on the right paw with a minimum 1-minute resting interval between each application, and the number of withdrawals was recorded. Mechanical sensitivity was expressed as % response frequency to each filament or as 60% response threshold (g), defined as the minimum pressure required for eliciting three out of five withdrawal responses (flinching, licking, or guarding the paw).

### Heat stimulation of mouse hindpaws (Hargreaves test)

Heat sensitivity was assessed by evaluating the hind paw withdrawal latency in response to radiant heat with the Hargreaves apparatus (Ugo Basile, Varese, Italy). Mice were kept in standard plastic modular enclosures on top of a glass platform (Ugo Basile, Varese, Italy) enabling the application of the radiant heat source (infrared intensity 40) to the hind paw plantar surface. Mice were acclimatized on three consecutive days for 1 h to the setup and at least during 30min before starting the measurements on each experimental day. Three measures were taken on each paw with a minimum 5 minutes resting interval between the stimulations and a cut-off time of 20s. The withdrawal latency was averaged for each animal’s paw on each day.

### Formalin test

The intraplantar formalin test was performed as described (Stösser et al., 2010): in brief, Formalin (5%, 20μl) was injected into the plantar surface of one hind paw, and the duration of nocifensive behaviors including lifting, licking, or flinching of the injected paw was measured in 5min bins for a duration of 50min after injection.

### Morris Water Maze

A standard hidden platform protocol was employed. The circular pool (r = 85 cm) was filled with opaque water to a height of 35 cm and a circular escape platform (r= 5 cm) was submerged 1cm below the water surface at a constant position in the center of the North-West (NW) quadrant during training. The pool openly faced the testing room which provided ample distal cues for visual spatial navigation. To reduce stress effects, mice were habituated to the maze 24 hours prior to training (4 trials, visible platform located once in every quadrant). Training consisted of 4 daily trials on 7 days split into 2 and 5 consecutive days with 2 days rest in between. Cages were placed under infra-red lamps to prevent hypothermia. Animals were introduced to the pool from start positions East (E), South-East (SE), South (S), and South-West (SW) to avoid close initial proximity to platform. Starting positions were block-randomized with all possible starting positions in random order on every day. After mounting the platform, animals were left there for 15 s. Cut-off time for trials were set to 60 s after which animals failing to locate the platform were guided with a wooden rod and left there for 15 s. One probe trial of 120 s with the platform removed from the pool was conducted 24 hours after the last training day. Here, animals entered the pool from position SE.

### Whole cell patch clamp

Mice were sacrificed by injecting 200μL of Ketamine/Xylacine (Ketamine: 220 mg/kg, Ketavet; Zoetis and Xylazine 16 mg/kg, Rompun; Bayer) in PBS and the extracted lumbar spinal cord portion was embedded in 2% low meting point agarose (Bio&SELL) and sliced with a vibratome (Leica VT1200S, Germany) into 300 μm sections using a slicing solution containing (in mM): sucrose, 191; K-gluconate, 0.75; KH2PO4, 1.25; NaHCO3, 3; D-glucose, 20; myo-inositol, 3; ascorbic acid, 1; choline bicarbonate, 23; ethyl pyruvate, 5; CaCl2, 1; MgSO4, 4. Slices were allowed to recover for 30 min at 32°C in recording aCSF containing (in mM): NaCl, 121; KCl, 3; NaH2PO4, 1.25; NaHCO3, 25; D-glucose, 15; ascorbic acid, 1; MgCl2, 1.1; myo-inositol, 3; CaCl2, 2.2; ethyl pyruvate, 5. Following recovery, slices were placed into the recording chamber and superfused at ∼2 ml/min with oxygenated recording aCSF. Whole-cell recordings from random lamina 1 neurons were performed using a patch-clamp amplifier (MultiClamp 700B, Axon Instruments, Molecular Devices, USA) and online data acquisition was performed with pClamp 11 (Axon Instruments, USA). The data were low-pass filtered at 10 kHz and sampled at a rate of 20 kHz. Electrodes (4–8 MΩ) were pulled from borosilicate glass capillaries (O.D. 1.5 mm, I.D. 0.86 mm, Sutter Instruments, USA). Intrinsic firing properties of lamina 1 neurons were recorded in the presence of 10 mM CNQX, 50 mM AP-V and 5 mM gabazine (Hello Bio) and using an internal solution containing (in mM): K-gluconate, 120; HEPES, 40; MgCl2, 5; Na2ATP, 2; NaGTP, 0.3. In some experiments aCSF was additionally supplemented with α-cyano-4- hydroxycinnamate (4-CIN, Sigma) or L-lactate (Sigma). In naïve animals, recorded neurons were randomly sampled from both dorsal horns; in CFA-injected animals, only the ipsilateral dorsal horn was sampled. Series resistance (Rs) was typically 10–30 MΩ across experiments. In current clamp recordings, pipette capacitance compensation and bridge balance were applied before the recording protocol. Current steps to record rheobase and firing properties were applied for 500 ms from 0 to 120 pA and at 2.5 pA intervals. The liquid junction potential between external and internal solutions of ∼12.8 mV was not corrected for.

### Seahorse Assay

#### Sample preparation

3-5 week old mice were sacrificed by Ketamine/Xylacine injection and the extracted lumbar spinal cord portion was embedded in 2% low meting point agarose (Bio&SELL) and sliced with a vibratome (Leica VT1200S, Germany) into 220 μm sections using a slicing solution continuously oxygenated containing (in mM): sucrose, 191; K-gluconate, 0.75; KH2PO4, 1.25; NaHCO3, 3; D-glucose, 20; myo-inositol, 3; ascorbic acid, 1; choline bicarbonate, 23; ethyl pyruvate, 5; CaCl2, 1; MgSO4, 4. Seahorse Assay in spinal cord tissue was adapted from a previously described protocol (Underwood et al., 2020) with some modifications. Briefly, slices were transferred to a holding chamber containing continuously oxygenated artificial cerebrospinal fluid (aCSF; 120 mM NaCl, 3.5 mM KCl, 1.3 mM CaCl2, 1 mM MgCl2, 0.4 mM KH2PO4, 5 mM HEPES, and 10 mM D-glucose; pH 7.4) and allowed to recover for 30 min at room temperature. Sections were individually transferred to a biopsy chamber containing fresh oxygenated aCSF. Rapid-Core biopsy & sampling punch with plunger system (500 µm; Micro-to-Nano, Netherlands) were used to take a sample of the dorsal horn. For naïve animals, samples were taken from both dorsal horns. In CFA-injected animals, only the Ipsilateral dorsal horn was sampled. Punches were ejected directly into an XFe96 Cell Culture Microplate (Agilent Seahorse XF, XF96 FluxPack) based on a pre-determined plate layout. Each well contained 180 µL room temperature assay media (aCSF supplemented with 0.6 mM pyruvate and 4 mg/ml BSA). After loading all biopsy samples, each well was visually inspected to ensure that the punches were submerged and centered at the bottom of the well. The XFe96 Cell Culture Microplate was then incubated at 37°C in a non-CO2 incubator for approximately 30 min. During this incubation period, 10x concentration of assay drugs (prepared in aCSF) were loaded into their respective injection ports of a XFe96 Extracellular Flux Assay sensor cartridge (previously hydrated for 24h with Agilent Seahorse XF Calibrant solution at 37°C in a non-CO2 incubator). The sensor cartridge containing the study drugs was then inserted into the analyzer for calibration. Once the analyzer was calibrated, the calibration plate was replaced by the microplate containing the tissue punches and the assay protocol initiated.

#### Mitochondrial respiration and Glycolysis Assay

Mitochondrial respiration and glycolysis assay were measured using the Seahorse Bioanalyzer (Agilent Seahorse, XF96 Bioanalyzer; Pike Winer & Wu, 2014)

For the mitochondrial respiration, oligomycin (10 µM final), FCCP (15 μM final), and a combination of rotenone and antimycin-A (20 mM and 10 mM final, respectively). The bioanalyzer was calibrated and the assay was performed using Mito Stress Test protocol as suggested by the manufacturer (Agilent Seahorse Bioscience,). The assay was run in one plate with 5-10 replicates per condition.

For the glycolysis assay, the same aCSF media as before was used, with the exception that it did not contain Glucose and it was supplemented with 2 mM glutamine and 1 mM sodium pyruvate. Injections of glucose with KCl (10 mM final and 25 mM final, respectively), oligomycin (10 μM final) and 2-deoxy-d-glucose (2-DG; 50 mM final) were diluted in the Agilent Seahorse XF Assay Medium and loaded onto ports A, B and C respectively. The bioanalyzer was calibrated and the assay was performed using Glycolytic Stress Test protocol as suggested by the manufacturer (Agilent Seahorse Bioscience,). The assay was run in one plate with 5-10 replicates per condition Seahorse Wave software was used to analyze metabolic data generated from both assays. The data from each assay was normalized to the total protein content with a Pierce™ BCA Protein Assay Kit following the manufacturer’s instructions.

### Statistical analysis

Data analysis: The sequencing reads were aligned by EMBL Genecore to mouse genome Mm10. FastQC was used for quality check and the differential expression analysis was performed with R following the Bioconductor RNA-Seq workflow developed by Love et al., 2015. The following packages were used during the data analysis: DESeq2, Rsamtools, GenomicFeatures, TxDb.Mmusculus.UCSC.mm10.ensGene, GenomicAlignments, AnnotationDbi, org.Mm.eg.db. Data was analyzed using R version for Linux 3.6.1 (https://www.r-project.org), RStudio for Linux version 1.1.463, RStudio for Windows version 4.1.2 and Matlab for Windows version R2016a- 2020a (MathWorks, USA). Statistical tests were performed using GraphPad Prism for Windows V7.00-8.0.1 (GraphPad software, USA) or RStudio for Windows version 4.1.2. Results are presented as mean ± standard error of the mean (SEM) unless indicated otherwise. Distribution of data was assayed using the KS normality test, the D’Agostino and Pearson omnibus normality test and the Shapiro-Wilk normality test. For statistical testing of data with only two groups, 2-tailed Student’s t-test was used. In case there were more than two groups with only one source of variation a One-way ANOVA followed by Tukey’s multiple comparison test was used. When comparing two or more groups with more than 1 source of variation, a Two-way ANOVA followed by Bonferroni or Sidak’s post hoc test. Star, ‘*’, signifies *P*<0.05, ‘**’ *P*<0.01, ‘***’ *P*<0.005, ‘****’ *P*<0.001.

### Data availability

Individual data points are represented throughout. More detailed datasets are available from the corresponding author upon reasonable request.

## Acknowledgements

We thank Bal Khakh for Aldh1L1-Cre^ERT2^ mice, Inaam Nakchbandi and Reinhard Fässler for the germline deleter Cre mouse line; Amandine Cavaroc, Annika von Seggern and Daniela Pimonov for expert technical support; Daria Kocherhina for help with Morris Water Maze tests; Erica Underwood for help with spinal slice Seahorse experiments; Ruth Drdla-Schutting and Maria Kronschläger for advice on spinal electrophysiology; members of the Siemens lab for inspiring discussions and critical input; Vladimir Benes and the EMBL genomics core facility for support with RNA sequencing and analysis; Claudia Pitzer and the Interdisciplinary Neurobehavioral Core (INBC) as well as The Nikon Imaging Center at Heidelberg University for support with confocal microscopy. The authors gratefully acknowledge the data storage service SDS@hd supported by the Ministry of Science, Research and the Arts Baden-Württemberg (MWK) and the German Research Foundation (DFG) through grant INST 35/1314-1 FUGG. This work was supported by the German research Foundation SFB1158 (to K.S.-S., A.T.-T., M.S., R.K. and J.S.; projects S02, S01, A07, A09, B01 and B06), the European Research Council ERC-CoG-772395 (to J.S.) and the International Human Frontier Science Program Organization postdoctoral fellowship LT000762/2019-L (to W.A.).

## Author contributions

J.S. together with S.M.-L. and S.L conceived the project. S.L. carried out the ribosomal capture screen, identified PTG and performed glycogen analysis together with S.M.-L. and with help from H.W. A.A.D.-R. generated conditional PTG^(-/-)^ mice. S.M.-L. characterized PTG^(-/-)^ mice, performed metabolic analysis together with T.F. and performed mouse pain behavior experiments with help from M.S., A.T.-T. and supervision and critical input from R.K. W.A. carried out all electrophysiological recordings. A.M.H. helped with spinal slice preparations and K.S.S and T.F. supported analysis and presentation of the data. J.S. wrote the paper. All authors commented on and approved the paper.

