## Extended Data for "Neuron-astrocyte metabolic coupling facilitates spinal plasticity and maintenance of persistent pain"

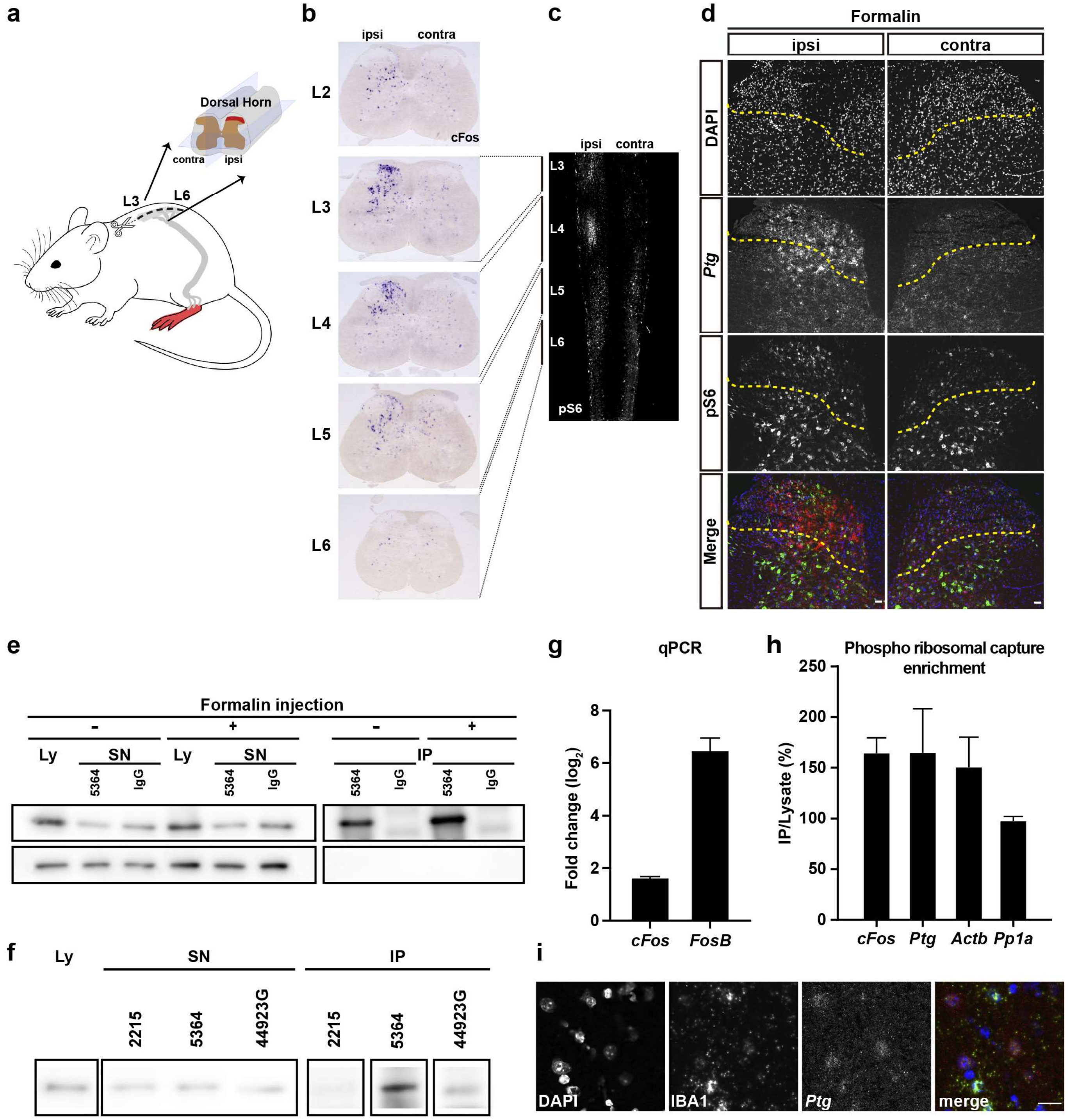

a
Dorsal Horn
contra
ipsi
L3
L6
b
ipsi
contra
L2
L3
L4
L5
L6
cFos
c
ipsi
contra
L3
L4
L5
L6
pS6
d
Formalin
ipsi
contra
DAPI
Ptg
pS6
Merge
e
Formalin injection
-
+
Ly
SN
5364
IgG
IP
g
qPCR
Fold change (log2)
cFos
FosB
h
Phospho ribosomal capture enrichment
IP/Lysate (%)
cFos
Ptg
Actb
Pp1a
f
Ly
SN
IP
2215
5364
44923G
i
DAPI
IBA1
Ptg
merge

**Extended Data Fig. 1: Discovery of *Ptg* as a pain-induced gene in the dorsal spinal cord, related to Fig. 1.**

**(a)** Cartoon illustrating the anatomy of the spinal cord and the tissue isolation of the ipsilateral (pain-affected) dorsal horn (red labeled) and the contralateral (unaffected control) dorsal horn of lumbar sections L3 to L6 for phospho-ribosome capture experiments after formalin injection into the left hindpaw.

**(b)** c-Fos mRNA expression (*in situ* hybridization) in the dorsal spinal cord 2 hours (h) after intraplantar formalin injection. *C-Fos* expression is specifically detectable on the ipsilateral side (side of injection) of lumbar sections L2 to 6.

**(c)** Spatial distribution of pS6 (whole mount immunofluorescence) along the spinal cord, 2h after intraplantar formalin injection. Maximal projection image of z-stack images of the dorsal horn shown.

**(d)** Representative images of combined immunohistochemistry and fluorescent *in situ* hybridization with an antibody directed against pS6 (green) and an *in situ* probe directed against *Ptg* (red), DAPI (blue); scale bar: 40μm.

**(e)** pS6 immunoprecipitation of ipsilateral (+) and contralateral (-) dorsal spinal cord 2h after formalin injection on the ipsilateral side. pS6 was detected in both ipsi- and contralateral lysates (Ly), reduced in supernatant (SN) and enriched in immunoprecipitation eluates (IP). IgG control did not bind any pS6. IP showed an enrichment for pS6 on the ipsilateral (+) over the contralateral (-) side.

**(f)** Western blot showing anti-pS6 antibody test (Cell Signaling polyclonal antibody #2215, Cell Signaling monoclonal antibody #5364 and Invitrogen polyclonal antibody #44923G) for pS6 immunoprecipitation using spinal cord tissue after formalin injection. Ly, the three SNs after antibody incubation and the respective IPs were loaded on one a western blot gel. All three tested antibodies reduced pS6 amount in the supernatant (SN) after immunoprecipitation while #5364 showed best enrichment in the IP.

**(g)** qPCR verification of cDNA libraries generated from phospho-ribosome captured ipsilateral and contralateral dorsal horn spinal cord tissue after formalin stimulation, amplifying *cFos* and *FosB* cDNAs. *cFos* and *FosB* showed significant and consistent enrichment in tissue isolated from the ipsilateral (formalin affected) side (N=3). Unpaired two-tailed t-test; data represent mean ± s.e.m. *$p < 0.05$.

**(h)** Comparison of phospho-ribosomal capture to purely lysate-extracted mRNA. qPCR was performed before and after phospho-ribosomal capture using ipsilateral and contralateral dorsal spinal cord tissue of mice injected with formalin on the ipsilateral side. Note that the indicated genes (except for *Pp1a*) were found to be enriched (showed a higher IP/Ly ratio) in the phospho-ribosomal capture (IP) condition compared to qPCRs done from ipsilateral tissue (Ly), demonstrating that phospho-ribosomal capture is more sensitive, albeit only slightly, in pulling out induced mRNA transcripts. Unpaired two-tailed t-test; data represent mean ± s.e.m. *$p < 0.05$.

**(i)** Representative images of combined immunohistochemistry and fluorescent *in situ* hybridization with an antibody directed against microglia marker IBA1 (green) and an *in situ* probe directed against *Ptg* (red), DAPI (blue); scale bar: 20μm.

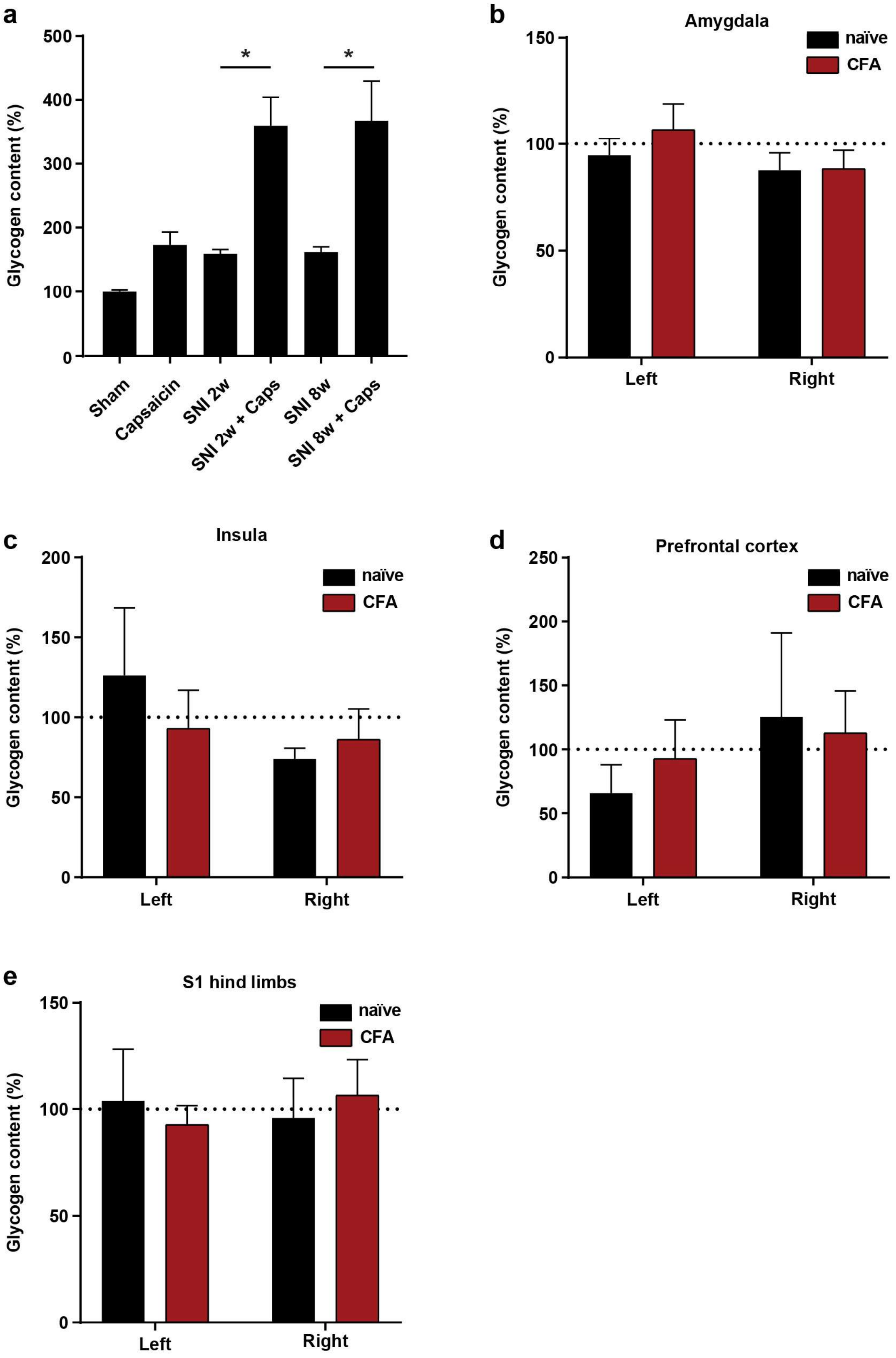
a
Glycogen content (%)
500
400
300
200
100
0
*
*
Sham
Capsaicin
SNI 2w
SNI 2w + Caps
SNI 8w
SNI 8w + Caps
b
Amygdala
naïve
CFA
150
100
50
0
Glycogen content (%)
Left
Right
c
Insula
naïve
CFA
200
150
100
50
0
Glycogen content (%)
Left
Right
d
Prefrontal cortex
naïve
CFA
250
200
150
100
50
0
Glycogen content (%)
Left
Right
e
S1 hind limbs
naïve
CFA
150
100
50
0
Glycogen content (%)
Left
Right

**Extended Data Fig. 2: Different to the spinal cord, pain stimuli do not promote major glycogen accumulation in higher brain centers processing pain signals, related to Fig. 2.**

**(a)** Glycogen content of ipsilateral dorsal spinal cord tissue compared to that of the contralateral side 2 or 8 weeks after Spared Nerve Injury- (SNI-)induced pain and with or without intraplantar capsaicin injection 6h prior to tissue preparation; expressed as percent glycogen content of the ipsilateral dorsal spinal cord of sham/untreated mice. Note that SNI or capsaicin alone only slightly elevate glycogen levels while the combination of the two synergistically increases glycogen accumulation. (N=3 mice); one-way ANOVA with Tukey's post hoc test.

**(b-e)** Glycogen content in the amygdala (**b**), insula cortex (**c**), prefrontal cortex (**d**) and hindlimb region of the somatosensory cortex (**e**) isolated from naïve or 6h CFA-treated mice. Glycogen content of these bilateral brain regions is expressed as percent of the average of the left and right side of the naïve mice. Two-way ANOVA with Bonferroni post hoc test; data shown as mean ± s.e.m. $*p < 0.05$.

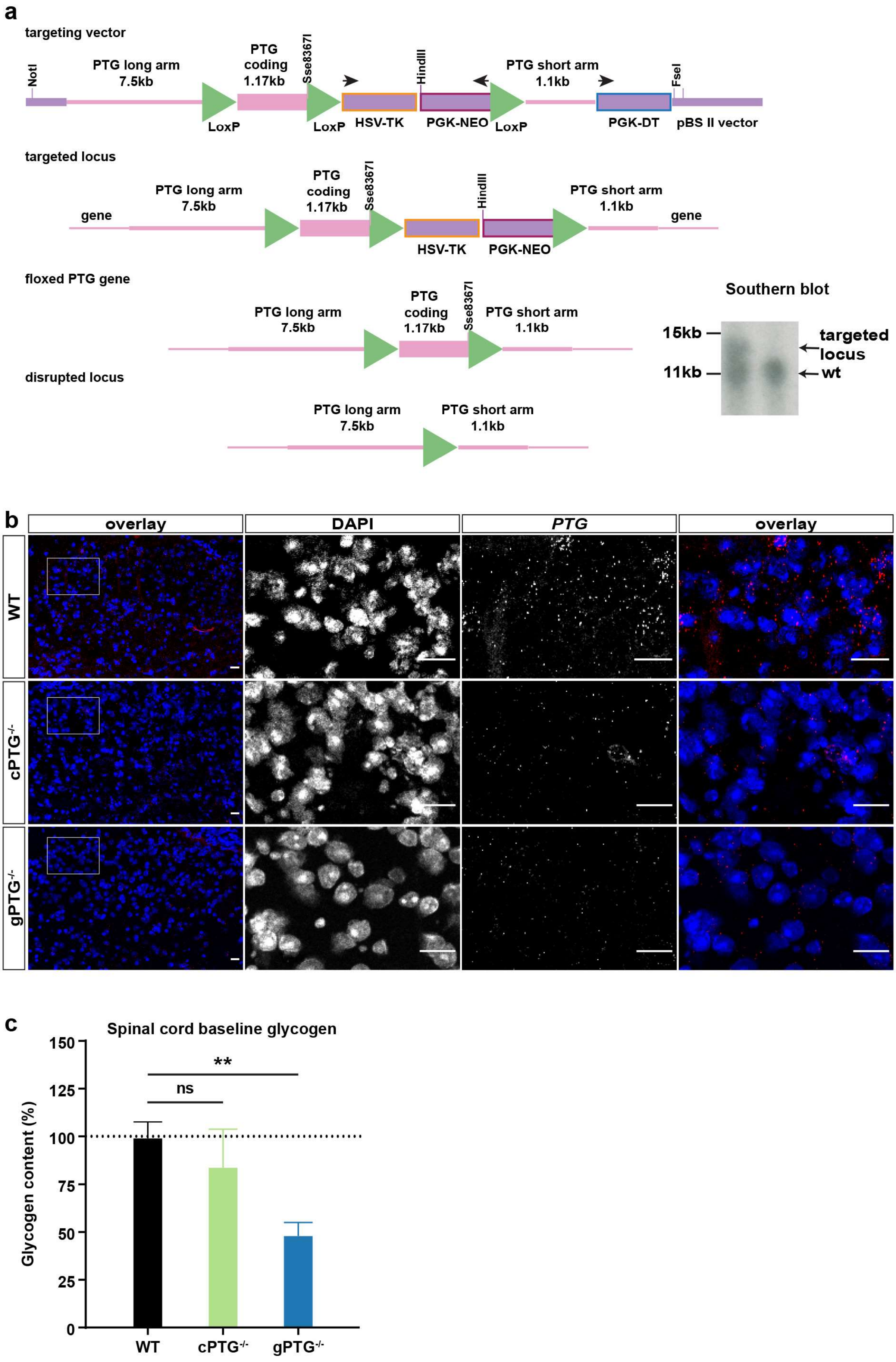
a
targeting vector
NotI
PTG long arm
7.5kb
PTG
coding
1.17kb
Sse8367I
HindIII
PTG short arm
1.1kb
FseI
LoxP
LoxP
HSV-TK
PGK-NEO
LoxP
PGK-DT
pBS II vector
targeted locus
gene
PTG long arm
7.5kb
PTG
coding
1.17kb
Sse8367I
HindIII
PTG short arm
1.1kb
gene
HSV-TK
PGK-NEO
floxed PTG gene
PTG long arm
7.5kb
PTG
coding
1.17kb
Sse8367I
PTG short arm
1.1kb
disrupted locus
PTG long arm
7.5kb
PTG short arm
1.1kb
Southern blot
15kb
11kb
targeted
locus
wt
b
overlay
DAPI
PTG
overlay
WT
cPTG-/-
gPTG-/-
c
Spinal cord baseline glycogen
Glycogen content (%)
150
125
100
75
50
25
0
**
ns
WT
cPTG-/-
gPTG-/-

**Extended Data Fig. 3: *Ptg* deletion blunts pain-induced glycogen build-up in the dorsal spinal cord, related to Fig. 3.**

**(a)** Embryonic stem cell targeting strategy to conditionally delete the single coding *Ptg* exon by introducing upstream and downstream loxP sites (green triangles). On the right panel a representative southern blot is shown, indicating the targeted locus by digesting genomic DNA with NcoI that cuts outside the two targeting arms resulting in a wildtype band of 11.5kb and a targeted band of 13kb.

**(b)** Representative images of *in situ* hybridizations (RNAscope) with a probe for *Ptg* (red) of ipsilateral dorsal spinal cord tissue from wildtype, $cPTG^{(-/-)}$ and $gPTG^{(-/-)}$ mice 2 hours after CFA stimulation. DAPI (blue), scale bar: 20 µm.

**(c)** Glycogen content (percent of WT) of dorsal spinal cord tissue from the three indicated mouse lines without any stimulation (WT: N=11; $cPTG^{(-/-)}$:N=5; $gPTG^{(-/-)}$: N=7); one way ANOVA with Tukey's post hoc test; data represent mean ± s.e.m. **p < 0.01.

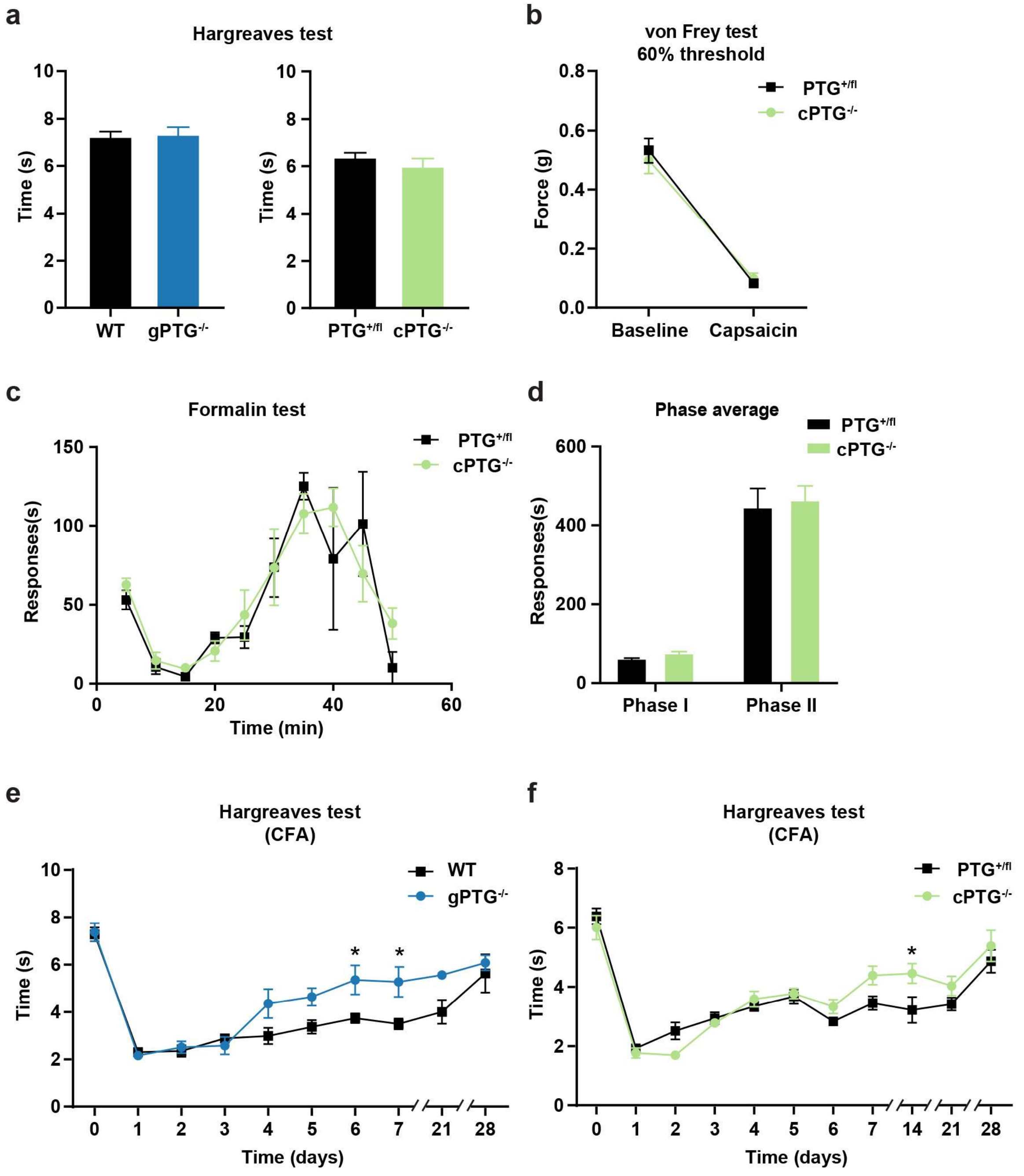
a
Hargreaves test
Time (s)
WT
gPTG-/-
PTG+/fl
cPTG-/-
b
von Frey test
60% threshold
Force (g)
Baseline
Capsaicin
c
Formalin test
Responses(s)
Time (min)
d
Phase average
Phase I
Phase II
e
Hargreaves test
(CFA)
Time (days)
f
Hargreaves test
(CFA)
Time (days)

**Extended Data Fig. 4: Acute pain is largely unperturbed in PTG$^{(-/-)}$ animals, related to Fig. 4.**

**(a)** Acute nocifensive responses upon heat stimulation using a focal light source directed onto the hindpaw of mice (Hargreaves test). Response latency was similar in cPTG$^{(-/-)}$, gPTG$^{(-/-)}$ and their respective wildtype or cPTG$^{(+/fl)}$ control mice (WT: N=6; gPTG$^{(-/-)}$, PTG$^{(+/fl)}$: N=12; cPTG$^{(-/-)}$: N=12); unpaired two-tailed t-test.

**(b)** Mechanical threshold required to elicit a response in at least 60% of trials in PTG$^{(+/fl)}$ and cPTG$^{(-/-)}$ mice that are either not treated (baseline) or 15 minutes after intraplantar capsaicin injection into the hindpaw (N=6); two-way ANOVA with Bonferroni post hoc test.

**(c, d)** Time course of formalin-induced nocifensive responses scored in 5 min bins (**c**) or separated in Phase I (0-10min) and Phase II (10-50min) (**d**) for cPTG$^{(-/-)}$ mice (N=4) and cPTG$^{(+/fl)}$ control mice (N=7). Two-way ANOVA with Bonferroni post hoc test.

**(e, f)** Nocifensive responses upon heat stimulation using a focal light source directed onto the hindpaw (Hargreaves test) measured before and at different time points after CFA-induced pain comparing WT (N=5) and gPTG$^{(-/-)}$ mice (E, N=5) and PTG$^{(+/fl)}$ and cPTG$^{(-/-)}$ mice (F, N=12). Two-way ANOVA with Bonferroni post hoc test; data represent mean ± s.e.m. *$p < 0.05$.

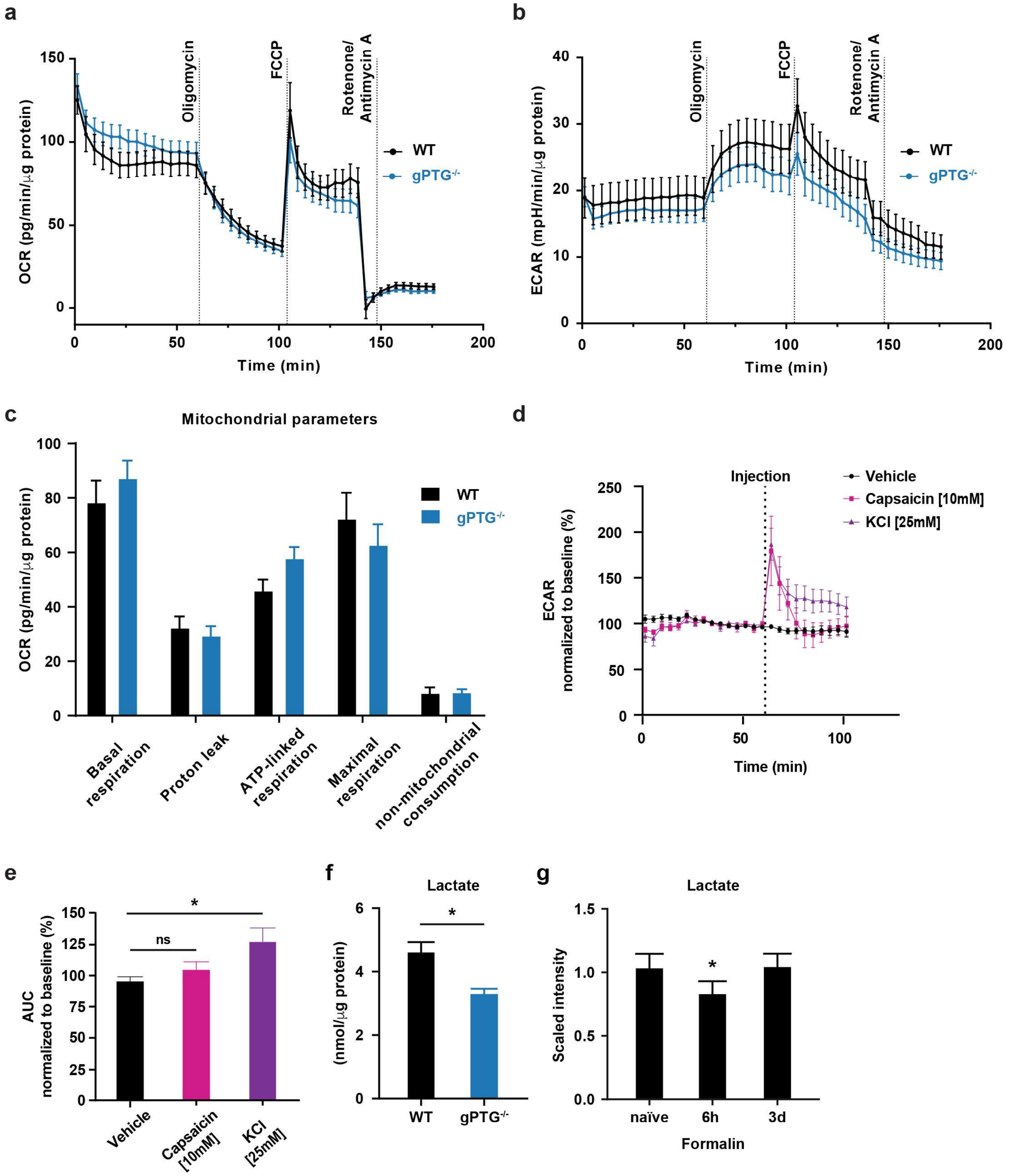
a
b
c
d
e
f
g
OCR (pg/min/μg protein)
ECAR (mpH/min/μg protein)
Time (min)
Oligomycin
FCCP
Rotenone/ Antimycin A
WT
gPTG-/-
Mitochondrial parameters
Basal respiration
Proton leak
ATP-linked respiration
Maximal respiration
non-mitochondrial consumption
Injection
Vehicle
Capsaicin [10mM]
KCl [25mM]
ECAR normalized to baseline (%)
AUC normalized to baseline (%)
ns
*
Lactate
(nmol/μg protein)
Scaled intensity
naïve
6h
3d
Formalin

**Extended Data Fig. 5: Basic spinal respiratory parameters are not perturbed in the absence of PTG, related to Fig. 5.**

**(a-c)** Seahorse mitochondrial-stress test in spinal cord dorsal horn slices comparing Oxygen Consumption Rate OCR **(a)**, Extracellular Acidification Rate ECAR **(b)** and mitochondrial parameters **(c)** between WT (n=37/5) and gPTG$^{(-/-)}$ (n=40/5).

**(d)** Effect of 10µM capsaicin and 25mM KCl on ECAR in WT spinal cord dorsal horn slices. Data normalized to the average baseline of each condition (n=7/1).

**(e)** Change of ECAR produced by the stimulus (capsaicin or KCl) measured as change of area under the curve (AUC) derived from data shown in (**d**; n=7/1).

**(f)** Lactate content in spinal cord dorsal horn from naïve WT and gPTG$^{(-/-)}$ mice. Unpaired two-tailed t-test (N=4).

**(g)** Lactate content after formalin pain stimulus, normalized to naïve animals and measured by mass-spectrometry (see methods). One-way ANOVA with Tukey's post hoc test (N=5). Data represent mean ± s.e.m. $*p < 0.05$.

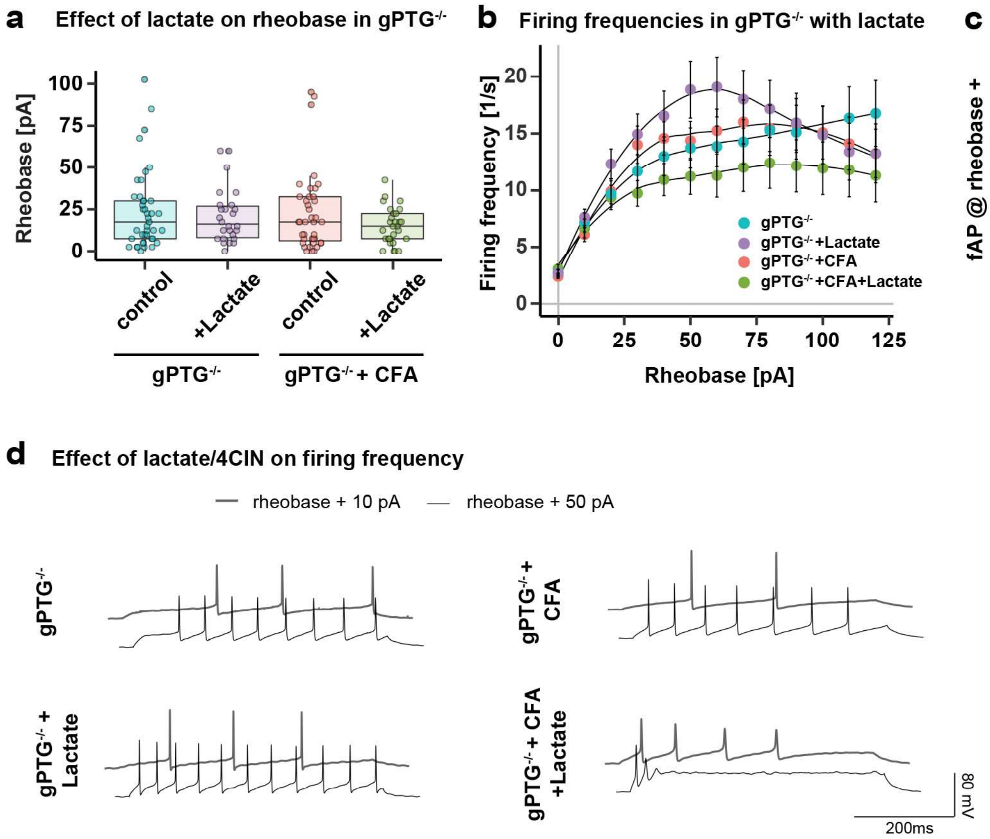

a
Effect of lactate on rheobase in gPTG-/-
Rheobase [pA]
100
75
50
25
0
control
+Lactate
control
+Lactate
gPTG-/-
gPTG-/- + CFA
b
Firing frequencies in gPTG-/- with lactate
Firing frequency [1/s]
20
15
10
5
0
0
25
50
75
100
125
Rheobase [pA]
gPTG-/-
gPTG-/- +Lactate
gPTG-/- +CFA
gPTG-/- +CFA+Lactate
c
fAP at +50 pA in gPTG-/-
fAP @ rheobase + 50 pA [Hz]
50
40
30
20
10
0
control
+Lactate
control
+Lactate
gPTG-/-
gPTG-/- + CFA
d
Effect of lactate/4CIN on firing frequency
rheobase + 10 pA
rheobase + 50 pA
gPTG-/-
gPTG-/- + CFA
gPTG-/- + Lactate
gPTG-/- + CFA +Lactate
80 mV
200ms

**Extended Data Fig. 6: Lactate is not sufficient to increase excitability of spinal L1 neurons in the absence of PTG, related to Fig. 6.**

**(a)** Rheobase comparison in $gPTG^{(-/-)}$/$gPTG^{(-/-)}$+CFA groups recorded with the addition of lactate (15mM) into recording aCSF. n=41 cells/N=4 animals ($gPTG^{(-/-)}$), n=26/3 ($gPTG^{(-/-)}$ + Lactate), n=39/4 ($gPTG^{(-/-)}$+CFA) and n=28/3 ($gPTG^{(-/-)}$+CFA+Lactate).

**(b)** Comparison of the firing frequencies in response to 500ms current injections (from 0pA to 120pA above rheobase) in naïve/CFA $gPTG^{(-/-)}$ groups in the presence of L-lactate.

**(c)** Comparison of firing frequencies at 50pA above rheobase current (based on **b**). n=41 cells/N=4 animals ($gPTG^{(-/-)}$), n=26/3 ($gPTG^{(-/-)}$ + Lactate), n=39/4 ($gPTG^{(-/-)}$+CFA) and n=28/3 ($gPTG^{(-/-)}$ +CFA+Lactate).

**(d)** Example traces of firing patterns of $gPTG^{(-/-)}$/$gPTG^{(-/-)}$+CFA neurons recorded in the presence of lactate. Data shown as mean ± s.e.m.

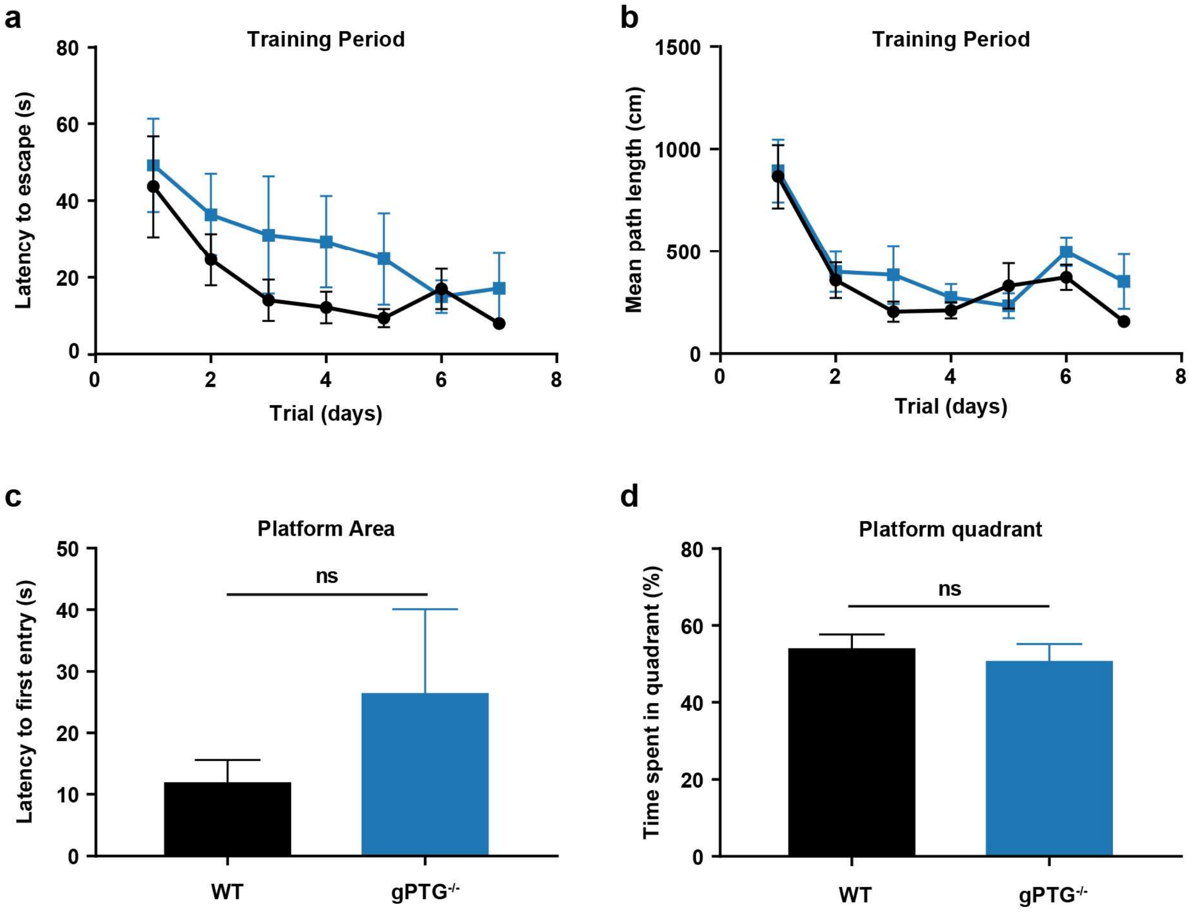
a
Training Period
Latency to escape (s)
80
60
40
20
0
0
2
4
6
8
Trial (days)
b
Training Period
Mean path length (cm)
1500
1000
500
0
0
2
4
6
8
Trial (days)
c
Platform Area
Latency to first entry (s)
50
40
30
20
10
0
ns
WT
gPTG-/-
d
Platform quadrant
Time spent in quadrant (%)
80
60
40
20
0
ns
WT
gPTG-/-

**Extended Data Fig. 7: Basic learning and memory is similar in wildtype and PTG$^{(-/-)}$ mice in the Morris Water Maze (MWM) test.**

**(a**, **b)** Latency to escape from swimming in the water and to reach a platform **(a)** and average distance traveled until the platform was reached **(b)** during the training period in the Morris Water Maze (MWM) of wildtype (WT) and gPTG$^{(-/-)}$ mice. Two-way ANOVA with Bonferroni post hoc test.

**(h**, **i)** Latency to first entry the platform area **(h)** and percent of time WT and gPTG$^{(-/-)}$ spent in the quadrant of the platform **(i)** during the last test of the MWM (8 days after start of training) when the platform has been removed. No differences in learning and memory where observed. Unpaired two-tailed t-test; N=16; data shown as mean ± s.e.m.
